# Correcting 4sU induced quantification bias in nucleotide conversion RNA-seq data

**DOI:** 10.1101/2023.04.21.537786

**Authors:** Kevin Berg, Manivel Lodha, Yilliam Cruz Garcia, Thomas Hennig, Elmar Wolf, Bhupesh K Prusty, Florian Erhard

## Abstract

Nucleoside analogues like 4-thiouridine (4sU) are used to metabolically label newly synthesized RNA. Chemical conversion of 4sU before sequencing induces T-to-C mismatches in reads sequenced from labelled RNA, allowing to obtain total and labelled RNA expression profiles from a single sequencing library. Cytotoxicity due to extended periods of labelling or high 4sU concentrations has been described, but the effects of extensive 4sU labelling on expression estimates from nucleotide conversion RNA-seq have not been studied. Here, we performed nucleotide conversion RNA-seq with escalating doses of 4sU with short-term labelling (1h) and over a progressive time course (up to 2h) in different cell lines. With high concentrations or at later time points, expression estimates were biased in an RNA half-life dependent manner. We show that bias arose by a combination of reduced mappability of reads carrying multiple conversions, and a global, unspecific underrepresentation of labelled RNA due to impaired reverse transcription efficiency and potentially global reduction of RNA synthesis. We developed a computational tool to rescue unmappable reads, which performed favourably compared to previous read mappers, and a statistical method, which could fully remove remaining bias. All methods developed here are freely available as part of our GRAND-SLAM pipeline and grandR package.

## INTRODUCTION

Nucleotide conversion sequencing of metabolically labelled RNA (1–3) enables the direct analysis of the temporal dynamics of RNA expression upon different perturbations in bulk or single cells (4–7) and of different quantitative parameters (8, 9). Cells are simultaneously subjected to a certain condition of interest and labelled with the nucleoside analogue 4-thiouridine (4sU), which is readily incorporated into nascent RNA. 4sU can be chemically converted in extracted RNA (1–3) or intact cells (7) to induce T-to-C mismatches in genome-mapped reads that originate from labelled RNA. Only a minor fraction (2-10%) of uridines is substituted by 4sU in nascent RNA, such that only a fraction of reads from labelled RNA molecules cover sites of 4sU incorporation (10). Statistical approaches nevertheless provide unbiased estimates of the new-to-total RNA ratio (NTR) and quantify the uncertainty in these estimates (11). Recent methods carry these uncertainties forward for the estimation of biophysical parameters of the temporal kinetics of RNA expression such as synthesis rates and half-lives and for identifying differentially regulated genes (12).

Labelling over several hours or using very high concentrations of 4sU affect cell viability (1) and rRNA processing (13). We recently showed that expression estimates from nucleotide conversion RNA-seq experiments can be affected before significant effects on cell viability are detectable (12). In different data sets, we observed marked differences between samples labelled with 4sU and unlabelled but otherwise biologically equivalent control samples in principal component analyses. In addition, in a differential gene expression analysis of total RNA levels, comparing 4sU labelled cells against equivalent unlabelled controls preferentially genes with short RNA half-lives appeared to be downregulated again. This observed downregulation can have biological or technical reasons (see Fig. 1): Excessive 4sU labelling might have direct effects on RNA metabolism, e.g. incorporation of 4sU into nascent RNA might result in reduced processivity of RNA polymerase II or 4sU containing RNA might be less stable than unlabelled RNA molecules. Excessive labelling might also induce indirect effects, e.g. due to the activation of cellular stress pathways.

**Figure 1.**
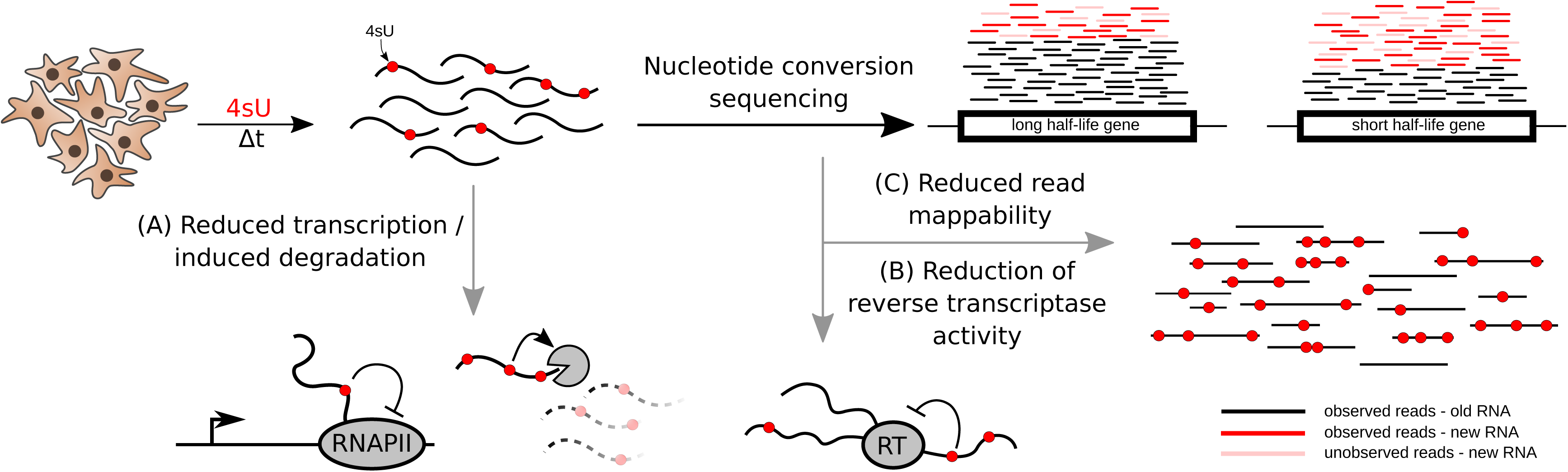
Overview of 4sU-induced quantification bias in new RNA. Cells are labeled with 4- thiouridine (4sU), which is incorporated into newly synthesized RNA. Incorporated 4sU could globally reduce transcriptional activity or induce degradation of labeled mRNAs (A) . 4sU has been shown to interfere with reverse transcription (B). T-to-C mismatches within read sequences makes it harder to correctly map reads (C). All three effects result in dropout of 4sU reads, mainly affecting genes with short half-lives and therefore introducing quantification bias (RNAPII: RNA polymerase II; RT: reverse transcriptase).

The observed downregulation of short-lived RNA might also have technical reasons apart from 4sU labelling before harvesting RNA for sequencing. In a recent study, reverse transcription efficiency was reduced for RNA containing 4sU converted with iodoacetamide (14) as it is used for SLAM-seq (1). The consequence of such a strong reduction of reverse transcription due to 4sU is that labelled RNA is underrepresented in the sequencing library for 4sU-labelled samples. Moreover, mismatched bases generally impact negatively on read mappability. If mappability is strongly impaired, reads corresponding to labelled RNA are underrepresented among the mapped reads used for quantifying gene expression. Since genes with short-lived RNAs have a higher percentage of labelled RNA in the total RNA pool than genes with long-lived RNAs, both, an underrepresentation of labelled RNA in the library, and an underrepresentation of labelled RNA in the mapped reads could explain quantification bias correlating with RNA half-lives.

Here, we performed nucleotide conversion RNA-seq with increasing concentrations of 4sU and with several periods of labelling in different cell types. We used these data to study the cell type specificity and dependence on the duration of labelling and 4sU concentration of biased expression estimates due to 4sU labelling and to assess the impact of technical reasons thereof. To counter these effects, we here propose a new method to rescue previously unmappable reads. We compared it to existing read mapping tools and evaluated it using in-silico simulated and real data sets. Furthermore, we devised a scaling strategy to correct for the underrepresentation of new RNA in the sequencing library or among mapped reads. Our data provides evidence that this correction completely removed this effect from the data enabling the analysis of samples that otherwise suffer from quantification bias due to excessive 4sU treatment.

## MATERIALS & METHODS

### Cell culture and 4sU labelling

NIH-3T3 (ATCC CRL-1658) Swiss murine embryonic fibroblasts, human U2OS (RRID:CVCL_0042), HFF-TERT (ATCC CRL-4001) hTERT-immortalized human foreskin fibroblasts and HCT 116 (RRID:CVCL_0291) cells were grown in DMEM (Dulbecco’s Modified Eagle’s Medium) supplemented with 100 IU/mL Penicillin and 100 mg/mL Streptomycin at 37°C/5% CO_2_. NIH-3T3 cells were supplemented with 10% NCS (New-born calf serum), U2OS, HFF-TerT and HCT116 cells were supplemented with 10% FBS.

All cells were seeded in six-well plates at 5*10 cells/well followed by 4sU-labelling the next day. NIH- 3T3 cells were labelled with 800 µM 4sU for 15, 30, 60, 90 and 120 minutes. U20S, HFF-TerT and HCT116 cells were labelled with 0, 100, 200, 400 or 800 µM 4sU for 1h.

The cell lines were routinely checked for Mycoplasma by PCR and tested negative at all times.

### SLAM-seq

Cells subject to 4sU-labelling were harvested using TRI reagent (Sigma) and RNA isolation was conducted using the Zymo DirectZol RNA-microprep kit (R2062) as described by the manufacturer and re-suspended in 1X PBS buffer. SLAM-seq (1) was conducted as described before (15) using IAA (Iodoacetamide) to mediate U>C conversions at 50 C/20 minutes. The reaction was quenched using excess DTT (Dithiothreitol). RNA was then purified using RNeasy Mini elute kit (Qiagen) and subject to quality control via gel electrophoresis for 18S and 28S RNA followed by Bioanalyzer assessment (Agilent 2100). Library preparation (Illumina TruSeq) and sequencing (2x75 pair-ended) was conducted by the Core Unit SysMed (Würzburg) using NextSeq500 as described previously (15).

### RNA-seq/SLAM-seq data processing

All publicly available RNA-seq and newly generated SLAM-seq data used here were processed using the GRAND-SLAM pipeline (11). Fastq files of publicly available RNA-seq data were downloaded from the SRA database. The accession numbers were: GSE162264 for the simulation of mismatches on read mappability and the evaluation of read mapping tools from Ref. 4 (sample: GSM4948135), GSE124167 (samples: GSM3523316- GSM3523318) and GSE109480 (Samples: GSM2944116 – GSM2944120) for the comparison of read mappability after T>C mismatch introduction in TruSeq (16) and QuantSeq (17) data sets respectively.

Adapter sequences were trimmed using cutadapt (version 3.5) using parameters “-a AGATCGGAAGAGCACACGTCTGAACTCCAGTCA -A AGATCGGAAGAGCGTCGTGTAGGGAAAGAGTGT” for the increasing concentrations and progressive labelling data and “-a AGATCGGAAGAGCACACGTCTGAACTCCAGTCA” for data from Ref (4). Then, bowtie2 (version 2.3.0) was used to map read against an rRNA (NR_046233.2 for TruSeq and QuantSeq data, and U13369.1 for increasing concentrations, progressive labelling and Ref. (4)) and Mycoplasma database using default parameters. Remaining reads were mapped against target databases using STAR (version 2.7.10b) using parameters “--outFilterMismatchNmax 20 --outFilterScoreMinOverLread 0.4 -- outFilterMatchNminOverLread 0.4 --alignEndsType Extend5pOfReads12 --outSAMattributes nM MD NH –outSAMunmapped Within”. We used the murine genome for TruSeq and QuantSeq data, the human genome for increasing concentrations, progressive labelling and data from Ref. (4). All genome sequences were taken from the Ensembl database (version 90 for human, version 102 for mouse). Bam files for each data set were merged and converted into a CIT file using the GEDI toolkit 33 and then processed using GRAND-SLAM (version 2.0.7) with parameters “-trim5p 15 -modelall” to generate read counts and NTR values on the gene level, taking into account all reads that are compatible with at least one isoform of a gene. For the newly generated SLAM-seq data sets only genes with more than 200 reads in half of the samples were retained for the evaluation of 4sU dropout scaling.

### Incorporation frequency saturation curves

We assume that the incorporation frequency of 4sU only depends on the relative concentrations of (triphosphorylated) U and 4sU. Ignoring 4sU uptake into cells and all steps necessary to make 4sU available for transcription, the incorporation frequency p can be computed from the U concentration *c_U_* and 4sU concentration *c*_4sU_ as 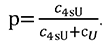. This p is shown as a function of *c*_4sU_ in Fig. 2D. The unknown U concentrations are estimated by solving this equation for 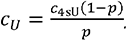, and taking the average of the *c_U_* computed from the 100μM and 200μM samples.

**Figure 2.**
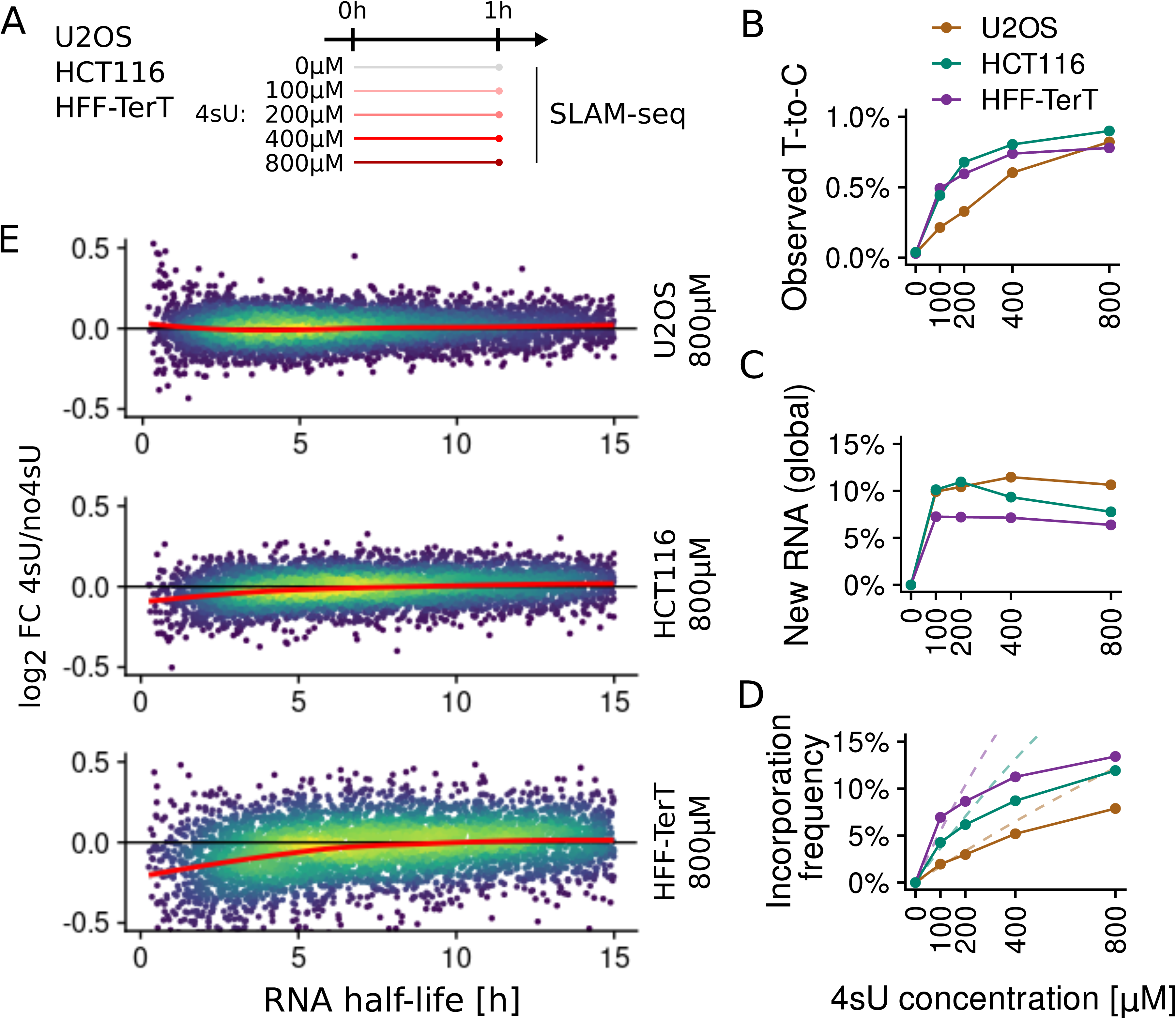
Evaluation of increasing concentrations on quantification bias. (A) SLAM-seq experiments were conducted with U20S, HCT116 and HFF-TerT cells labeled for 1h with 4sU concentrations of 0 µM, 100 µM, 200 µM, 400 µM or 800 µM. (B) Observed T-to-C mismatches across all reads per cell line and 4sU concentration. (C) Percentages of new RNA content per cell line and 4sU concentration estimated by GRAND-SLAM. (D) Incorporation frequency of 4sU into newly synthesized RNA per cell line and 4sU concentration estimated by GRAND-SLAM (solid line). The dashed lines represent the theoretically expected incorporation frequency (see Methods). (E) 4sU dropout plots of n=6,454 genes for the 800µM 4sU samples vs. 4sU naïve samples for all three cell lines. The x axis shows the RNA half-life, the y axis the median centered log2 fold change of total RNA expression for the 4sU labelled sample vs. the corresponding 4sU naïve control sample. A local polynomial regression (loess) fit is indicated in red.

### 4sU dropout plots

4sU dropout plots are computed for a sample labelled with 4sU and a biologically equivalent 4sU naïve control sample. The x axis of 4sU dropout plots is the RNA half-life computed from the 4sU labelled sample using the formula (11)

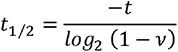

Here, *t* is the labelling time and v the new-to-total RNA ratio (NTR) estimated by GRAND-SLAM (11). Alternatively, the y axis is the NTR rank among all genes. The y axis is the log_2_ fold change of the labelled vs the naïve sample computed using the lfc package (18). These plots can be generated using the function Plot4sUDropout or Plot4sUDropoutRank of grandR (12).

### Simulation of nucleotide conversion RNA-seq with defined incorporation rates

We used the mapped reads from Ref. (4) to obtain the corresponding mapping positions and genes for each read sequence while unmapped reads were removed from the simulation. To classify reads as new or old, we used NTR values estimated from the data of Ref. (4) as follows: For a gene with a read count of *n* and an NTR value *v*, we randomly selected *n* · *v* reads and defined them as new.

In each new read, we randomly introduced additional T-to-C mismatches into the sequence by mutating a T in the read sequence into a C with probability equal to to a defined incorporation rate. Pre-existing T-to-C mismatches were kept. To simulate reads with shorter read lengths we trimmed the 3’ ends to the desired length.

Simulations based on the QuantSeq (17) and Illumina TruSeq (16) experiments where performed in the same manner to evaluate the effect of additional T-to-C mismatches on mappability in different library preparation methods and read lenghts.

To evaluate and compare the mapping accuracy of STAR, grand-Rescue, SLAM-DUNK, HISAT2 and HISAT-3N we generated read sequences fully in silico. We created 75 bp read sequences equal to the read counts per gene in the dataset from ref. (4) from random exonic locations of the respective genes. To simulate polymerase and technical errors, the reads were subjected to a 0.2% error rate by randomly altering single nucleotides. Then, T-to-C conversions were introduced as described above.

These reads were then mapped by all mapping tools, using the following parameters: For STAR we used “--outFilterMismatchNmax 20 --outFilterScoreMinOverLread 0.4 -- outFilterMatchNminOverLread 0.4 --alignEndsType Extend5pOfReads12 --outSAMattributes nM MD NH –outSAMunmapped Within”, for HISAT2 “--no-repeat-index”, for HISAT-3N “--base-change T,C -- no-repeat-index” and standard parameters for SLAM-DUNK.

### Pseudotranscriptome generation

As a basis for the creation of the pseudotranscriptomes of the homo sapiens and mus musculus genomes, we used the fasta- and gtf-files from the ensembl versions 90 and 102, respectively. For each genome, we processed the gtf-files to keep all entries with the gene, exon or CDS feature. Coordinates were change to reduce intronic and intergenic regions to 100 nucleotide spacers and genes on the negative strand were projected onto the plus strand. We then processed the fasta files, accordingly, removing intronic and intergenic sequences and replacing them by a uniform spacer of 100 N nucleotides. To transfer genes from the negative strand to the plus strand, their sequences were replaced by their reverse complements. Finally, all T nucleotides were exchanged by C.

### grand-Rescue

grand-Rescue is a two-step process, that starts from a bam file (containing read mappings without rescuing 4sU labelled reads) and generates a new bam file (additionally containing the rescued reads).

After mapping fastq files with STAR, grand-Rescue extracts unmapped reads, using the command “gedi -e ExtractReads” with standard parameters, writing all unmapped read sequences to a new fastq file, converting all T nucleotides to C and saving the original sequence per read along with the read IDs and all of its bam file tags to an idMap file. The resulting fastq file was then mapped to the pseudotranscriptome, using STAR with the following parameters: “--outFilterMismatchNmax 10 -- outFilterScoreMinOverLread 0.4 --outFilterMatchNminOverLread 0.4 --alignEndsType Extend5pOfReads12 --outSAMattributes nM MD NH –outSAMmode Full”. Afterwards, we removed all multimapped reads from this file with samtools (version 1.13) with the parameters “view -b -F 256”.

Subsequently, “gedi -e RescuePseudoReads” is used to transfer the mapping position to the original genome by using the mapped position in the pseudotranscriptome. We first identified the gene a read was mapped to in the pseudotranscriptome and the gene’s location on the plus or minus strand in the original genome. We calculate the distance of the alignment start position to the gene start position in the pseudotranscriptome and determine its alignment start position in the original genome by adding this distance to the gene start position in the original genome (or subtracting it from the gene end position, if the gene is originally on the negative strand and reverse complementing the sequence), skipping over intronic regions that may exist. Then, we recover the original read sequence before full T-to-C conversion along with all saved bam file tags and recalculate the nM and MD tags.

Finally, we remove unmapped reads from the original bam file and merge it with the rescued bam file from the pseudotranscriptome mapping using samtools.

### Correcting 4sU dropout

The percentage of 4sU dropout can be estimated for a sample labelled with 4sU if there is a biologically equivalent 4sU naïve control samples. It is estimated by numerically finding a factor *f* such that, if the NTR is multiplied by *f*, the spearman correlation coefficient of the log2 fold change 4sU/no4sU vs the NTR rank is 0. The 4sU dropout percentage then is 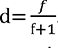. To correct for 4sU dropout, the expression of labelled RNA is multiplied by 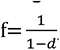, and the total expression estimate and the NTR is changed accordingly.

### RNA half-lives and testing for mis-normalization

RNA half-lives and 95% confidence intervals were estimated from the progressive labelling data using the non-linear least squares method described in Ref. using the grandR function FitKinetics after recalibrating effective labelling times using the grandR function

CalibrateEffectiveLabelingTimeKineticFit. The likelihood ratio test for an upward or downward trend in the total RNA of uncorrected data was performed by using the 4sU labelling time as independent variable in the target model, and only an intercept term for the background model. Testing was performed using the LikelihoodRatioTest function of grandR.

### Reverse transcription analysis

To determine the RNA fragments for the paired-end reads, we first used kallisto (version 0.44.0) with parameter --rf-stranded to infer transcript level expression for the two pooled 4sU naïve samples from our progressive labelling time course. We determined the major isoform for each gene by identifying the transcript with highest TPM value per gene, and the major isoform percentage by dividing the TPM of the major isoform by the total TPM of all transcripts for a gene. All genes with RNA half-life <30min, TPM>10 and a major isoform percentage of >90% were considered further. For each of these genes, and each sample, we collected all mapped read pairs, and determined the RNA fragment by connecting the two mates according to the exon-intron pattern of the major isoform. The corresponding sequences was used to count all k-mers with k=1…3.

## RESULTS

### Excessive 4sU treatment results in quantification bias preferentially for short-lived RNAs

We previously observed short-lived RNAs to be downregulated when comparing samples that were treated with 4sU for long periods of time (8h) to 4sU naïve samples (12). We reasoned that this could be caused by three effects (Fig 1): First, long-term treatment by 4sU could globally impact on RNA metabolism by reducing transcriptional activity or accelerating degradation of labelled RNA. With reduced transcriptional activity, all RNAs are inhibited by the same factor, but levels of short-lived RNAs would drop more rapidly than levels of long-lived RNAs, thereby explaining our observation. Second, as described previously (14), converted 4sU might reduce reverse transcription efficiency, such that labelled RNA is underrepresented in the sequencing library. The total number of T-to-C conversions for short-lived RNAs is larger than for long-lived RNAs, and, therefore, this could also explain the apparent downregulation of short-lived RNAs in 4sU treated samples. Third, the probability that a read is correctly mapped to its genomic locus of origin declines with increasing numbers of mismatched bases, which would also have its strongest effect on short-lived RNAs. Importantly, in all three cases fewer reads corresponding to newly synthesized RNA are mapped in the 4sU treated sample than in the 4sU naïve sample. In the first case, these reads are missing due to reduced RNA levels in the cells. In the second case, RNA levels are unaltered, but the composition of the sequencing library is changed. In the third case, the representation of genes by sequencing reads is unchanged but the read mapping algorithm could not assign them to their correct genomic loci.

To further investigate these, not mutually exclusive, causes, we generated several nucleotide conversion RNA-seq data sets of 4sU labelled samples: We performed dose escalation experiments by labelling with 0 μM, 100 μM, 200 μM, 400 μM and 800 μM of 4sU for 1h in the two cancer cell lines U2OS and HCT116 as well as in human telomerase reverse transcriptase immortalized primary foreskin fibroblasts (HFF-TerT; Fig. 2A). The observed 4sU-induced T-to-C conversions among all mapped reads increased with higher concentrations for all three cell lines (Fig. 2B). Interestingly, the maximal value at 800 μM was remarkably similar among the three cell lines (HFF-TerT, 0.80%; U2OS, 0.87%; HCT116, 0.94%), but the temporal kinetics were quite different. To investigate this further, we used GRAND-SLAM (11) to estimate the 4sU incorporation frequency (percentage of T-to-C conversions among labelled RNA only) and the percentage of labelled RNA for all samples. Except for HCT116, the estimated percentage of labelled RNA was largely constant for all concentrations indicating that GRAND-SLAM could reliably deconvolute the observed T-to-C conversions into contributions of the percentage of labelled RNA and different 4sU incorporation frequencies (Fig. 2C). Consistent with previous observations (19), incorporation frequencies among the three cell lines differed substantially and, as expected, increased with higher concentrations (Fig. 2D). The incorporation frequency only depends on the relative concentrations of activated 4sU and uridine (U) and is therefore expected to be approximately a linear function of the 4sU concentration in the regime well below the U concentration (see Methods). However, this increase saturated for all three cell lines well below 100%, indicating that, with increasing 4sU concentrations, import or activation of 4sU became rate limiting or that 4sU containing reads were underrepresented for the three reasons introduced above.

To investigate the effect of 4sU on short-lived RNAs we performed 4sU dropout analysis by correlating the log_2_ fold change of each 4sU treated sample to the corresponding 4sU naïve control sample of the same cell line vs the NTR. Interestingly, U2OS, which had the overall lowest incorporation frequencies (Fig. 2D), did not show downregulation of short-lived RNA even with 800 μM 4sU (Fig. 2E). By contrast, HCT116 and especially HFF-TerT showed this effect at higher concentrations (Fig. 2E).

We also sequenced a time course of murine NIH-3T3 fibroblasts labelled using 800 μM 4sU for 0 min, 15 min, 30 min, 60 min, 90 min and 120 min (Fig. 3A). Here, as expected, the raw T-to-C conversions as well as the percentage of newly synthesized RNA increased with longer periods of labelling (Fig. 3B-C). Interestingly, consistent with the observations made by us and others that activated 4sU accumulates only slowly in cells (12, 20), the incorporation frequencies of 4sU also increased from 2% for the 15 min and 30 min timepoints to more than 4% at the 1h and 2h time point (Fig. 3D). 4sU dropout analyses did not reveal any effects of 4sU up to 30 min treatment but showed increasingly stronger downregulation of short-lived RNAs at later time points (Fig. 3E).

**Figure 3.**
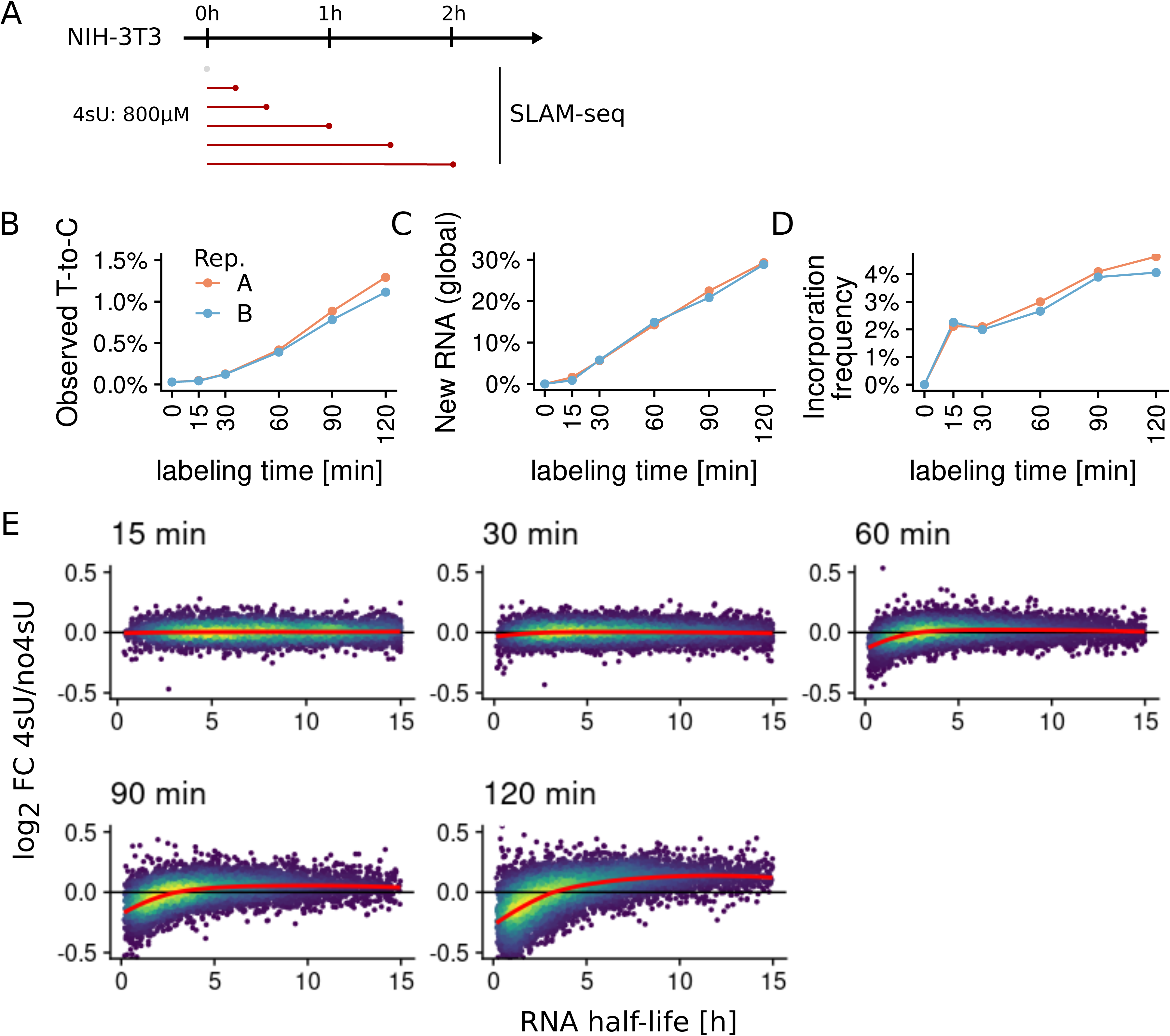
Evaluation of progressive labelling durations on quantification bias. (A) SLAM-seq experiments were conducted with NIH-3T3 cells with labelling for 0 min, 15 min, 30 min, 60 min, 90 min or 120 min with 800 µM of 4sU. (B) Observed T-to-C mismatches over all reads per time point and replicate. (C) Percentages of new RNA per time point and replicate estimated by GRAND-SLAM. (D) Incorporation frequency of 4sU into newly synthesized RNA per time point and replicate estimated by GRAND-SLAM. (E) 4sU dropout plots of n=9,072 genes for all time points of replicate A. The x axis shows the RNA half-life, the y axis the median centered log2 fold change of total RNA expression for the 4sU labelled sample vs. the corresponding 4sU naïve control sample. A local polynomial regression (loess) fit is indicated in red.

In summary, both increasing concentrations of 4sU as well as extended periods of labelling bias the quantification of total RNA due to less observed sequencing reads in short lived RNAs.

### Introduction of additional mismatches impairs read mappability

We first investigated whether the observed downregulation of short-lived RNA is solely due to diminished mappability of reads with T-to-C conversions. To this end, we considered all mapped reads (76bp, single-end) from a 4sU naïve control sample of a recent study (4) as a starting point and artificially and randomly introduced T-to-C conversions with varying incorporation frequencies ranging from 0% (equal to the original data) up to a maximum of 25% into the reads. This was done only for a fraction of the reads corresponding to the gene-wise NTR estimated in the original data, thus simulating a realistic nucleotide conversion sequencing experiment with controlled 4sU incorporation. We then used STAR (21) to map these reads back to the reference genome.

First, we investigated how many of the introduced T-to-C conversions were lost due to reduced mappability. As expected, higher incorporation frequencies directly correlated with the amount of lost T-to-C conversions, reaching already 9.8% at 10% incorporation rate and a maximum of 38.7% in the 25% sample (Fig. 4A). Interestingly, the number of lost mismatches did not rise linearly with increasing incorporation rates, indicating that reads with multiple mismatches are increasingly difficult to map. To test this, we binned the reads according to their number of introduced T-to-C conversions and analysed the count ratio of mappable reads vs the simulated reads in each bin. This ratio dropped steeply with every additional mismatch, resulting in a loss of 19% of reads with 3, and almost 50% of the reads with 5 T-to-C conversions (Fig. 4B). Thus, mappability suffers substantially in presence of multiple 4sU induced nucleotide conversions on the same read. Generally, mappability of reads decreases with increasing T-to-C conversion rates, but to a varying degree in relation to sequencing technique and read lengths (Fig. S1).

**Figure 4.**
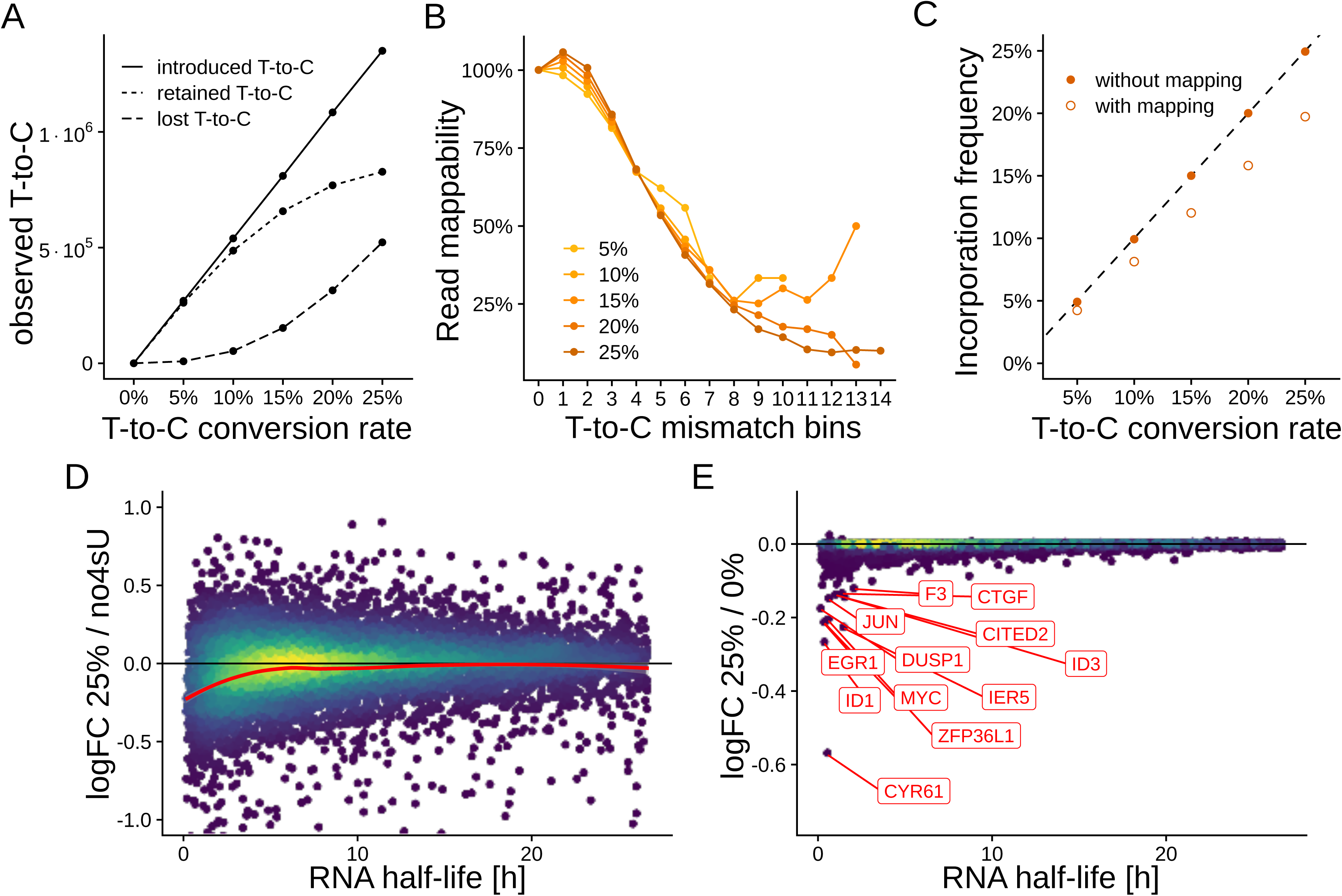
In-silico simulation of nucleotide-conversion RNA-seq (A) Line plots showing the number of introduced, retained and lost T-to-C mismatches after simulating T-to-C conversion rates of 0% up to 25% in 4sU naïve reads and subsequent remapping with STAR. (B) Line plots showing the percentage of all reads with 0 up to 14 T-to-C mismatches after mapping versus the true number of reads created by SLAM-seq simulations with 5% to 25% conversion rates. (C) GRAND-SLAM estimates of incorporation frequencies in newly synthesized RNA after introduction of T-to-C mismatches into read sequences and subsequent mapping (empty dots, with mapping) and into already mapped reads (solid dots, without mapping) for conversion rates of 5% to 25%. (D) 4sU dropout plot of a simulated 25% conversion rate sample vs. a 4sU naïve sample. (E) 4sU dropout plot of a simulated 25% conversion rate sample vs. the true read counts from the 0% conversion rate sample. All genes with a log2 fold change < -0.12 are highlighted.

Next, we used GRAND-SLAM to estimate incorporation frequencies in labelled RNA. Interestingly, the incorporation frequencies were underestimated by a fixed factor of approximately 0.83 for all simulated samples (Fig. 4C). This was an effect of read mapping, as introducing the mismatches into already mapped reads before running GRAND-SLAM resulted in unbiased estimates (Fig. 4C). Underestimation by a fixed factor is not unexpected since GRAND-SLAM utilizes the proportions of read counts with >1 T-to-C conversions for estimation of the incorporation frequency which suffer to the same extent from reduced mappability independent of the true incorporation frequency (Fig. 4B). The estimated incorporation frequency is an important parameter for the estimation of gene-wise NTRs. However, gene-wise NTR estimates were not biased due to underestimated incorporation frequencies of up to 15% (Fig. S2) and were underestimated specifically for high NTR values for incorporation frequencies above 15%. This indicates that the GRAND-SLAM model inherently compensates for biased estimates of incorporation frequencies when T-to-C mismatches are unobserved and only suffers when a substantial fraction of the reads is missing.

Finally, we compared our simulated samples with a 4sU naïve sample from the original data set to mimic 4sU dropout analyses of real data. Interestingly, similar to real data with high 4sU concentrations, short-lived RNAs appeared to be downregulated (Fig. 4D). However, this effect was only apparent at simulated incorporation frequencies of >15% and generally less pronounced as in the extreme cases of real data. We also compared the 25% sample with the 0% sample, which reflects the original reads without introduced T-to-C conversions, thereby removing variance between replicates. This revealed the loss of reads of short-lived key transcription factors like MYC and JUN or central signalling molecules like CYR61 (Fig. 4E). Gene set enrichment analysis revealed 199 gene ontology terms that consist of short-lived RNAs and therefore appear to be downregulated due to reduced mappability (Table S1). We concluded that albeit contributing, reduced mappability cannot explain the drastic loss of reads from short-lived RNAs observed in real samples treated with high concentrations of 4sU but might still result in biased fold changes for highly relevant classes of genes.

### grand-Rescue improves mappability of T-to-C conversion reads

Since reduced mappability of reads with T-to-C conversions has significant effects on quantification, we wondered whether read mapping could be improved. A promising approach that has been used in the past for other applications that involve nucleotide conversion such as bisulfite sequencing is to perform read mapping under a three-letter alphabet, e.g. after changing all T to C in both reads and reference (22). In principle, after switching to a three-letter alphabet, any standard read mapping tool can be used. Among the plethora of available read mapping tools, there are large differences in terms of mapping accuracy also without nucleotide conversion (23). We therefore favoured an approach that can use any available tool to do the actual read mapping, instead of adapting an existing tool or developing a new tool. Our method termed grand-Rescue is a two-step algorithm that first tries to map all reads to the reference genome without any modification, and then subsequently tries to map all unmappable reads to a pseudotranscriptome with a reduced alphabet. The final mapping locations are then transferred to the original reference genome (Fig. 5A). We use STAR (21) as the internal read mapper, which we found to have superior performance over several other read mapping tools.

**Figure 5.**
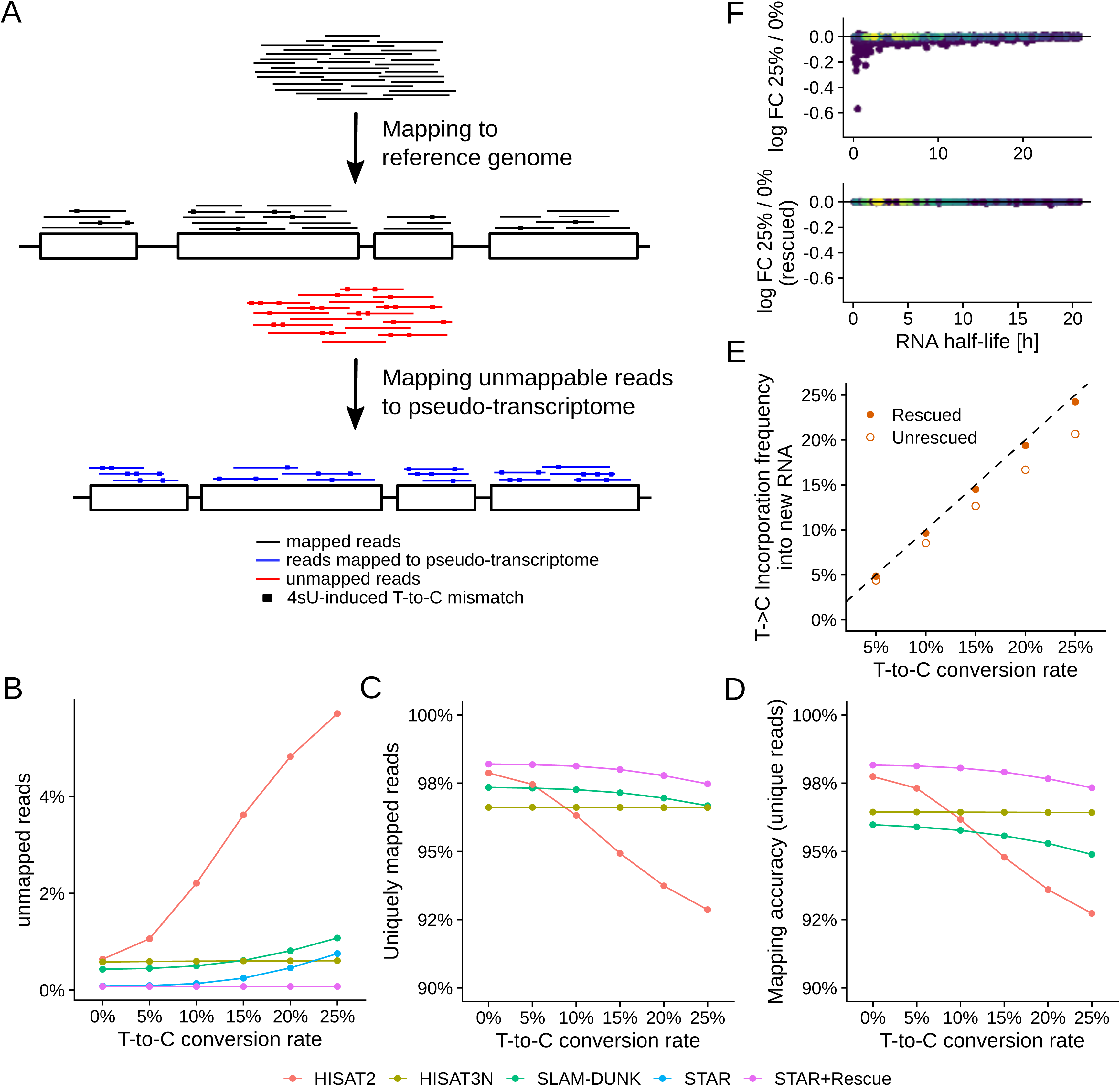
Evaluation of grandRescue and comparison to existing mapping tools(A) grandRescue first extracts unmappable reads, converts all T in their sequence to C and maps these reads with a read mapping tool of choice to a three-letter (T converted to C) pseudo-transcriptome. Rescued reads are then transferred to the original genome. (B) Percentage of unmapped reads for HISAT2, HISAT3N, SLAM-DUNK, STAR and STAR+Rescue for different simulated incorporation frequencies. (C) Percentage of uniquely mapped reads in relation to all reads per sample for HISAT2, HISAT3N, SLAM- DUNK, STAR and STAR+Rescue. (D) Correctly mapped reads in relation to all reads for HISAT2, HISAT3N, SLAM-DUNK, STAR and STAR+Rescue. (E) GRAND-SLAM estimates of incorporation frequencies before (empty dots) and after (solid dots) rescue. (F) 4sU dropout plots of a simulated 25% conversion rate sample vs. the true read counts from the 0% conversion rate sample before (top) and after (bottom) rescue.

We used simulated nucleotide conversion sequencing data to evaluate and compare the performance of grand-Rescue with STAR (21) and HISAT2 (24), two standard read mappers, as well as SLAM-DUNK (25) and HISAT3N (26), tools that have been developed specifically for nucleotide conversion RNA-seq. Instead of using STAR mapped reads as starting point as above, which would favour STAR based read mapping, we randomly redistributed reads across within their mRNA (see Methods). As expected, the percentage of unmappable reads for STAR and especially HISAT2 increased drastically with the conversion rate (Fig. 5B). Surprisingly, this was also the case for the T-to-C conversion aware read mapper SLAM-DUNK, while HISAT3N and grand-Rescue remained unaffected by increasing T-to-C conversions. Among the reads mappable by each individual tool, the percentage of uniquely mappable reads with increasing T-to-C conversions stayed constant for HISAT3N and only dropped slightly for both grand-Rescue and SLAM-DUNK (Fig. 5C). Importantly, however, HISAT3N and SLAM-DUNK only mapped 96.6% of the reads uniquely, while 97.8% where uniquely mapped by grand-Rescue even for the 25% sample. We observed a similar picture when analyzing the percentage of correctly mapped reads among unique reads, i.e. the mapping accuracy: For both, grand-Rescue and HISAT3N the accuracy did not drop with increasing T-to-C conversions, but HISAT3N had overall lower performance than grand-Rescue across all samples (Fig. 5D).

We concluded that all three T-to-C conversion aware read mappers, which follow different strategies, can indeed improve read mapping and that grand-Rescue performs favourably. Importantly, however, the internally used read mapping algorithm, which can be changed for grand-Rescue, also has a great effect on read mappability.

### grand-Rescue mitigates effects of reduced mappability

Rescuing previously unmappable T-to-C conversion reads using grand-Rescue substantially improved the estimates of the T-to-C conversion frequency which were now only slightly underestimated by a factor of roughly 0.97 instead of 0.83 before rescue (Fig. 5E). More importantly, after rescue, the apparent downregulation of short-lived RNA due to read mappability was not observed anymore (Fig. 5F). To account for the impact of different library preparation protocols and read lengths on the estimation of 4sU incorporation rates by GRAND-SLAM, we used different starting points for our simulation, including data generated using a 3’ end sequencing protocol (QuantSeq) as well as paired-end and single-end data sets based on random priming (TruSeq), and simulated 4sU incorporation rates from 0% to 10%. The QuantSeq data consisted of 75bp single end reads, whereas the TruSeq data were sequenced with 2x 125bp paired end reads. To mimic other sequencing modes, we in-silico trimmed the TruSeq reads to 100 or 75 bp reads and also discarded the second reads, and thus analyzed overall six settings based on TruSeq data, each with different incorporation rates (Fig. S3). The percentage of rescued reads was highest in QuantSeq, especially in the sample with a 10% conversion rate with 0.58% of all reads being rescued whereas in the single end TruSeq samples, less reads were rescued (Fig. S3A). These findings are also reflected in the incorporation estimation, which was underestimated most in 75 bp in QuantSeq and could be rescued (Fig. S3B) and to a minor extent in the TruSeq data sets, whereas longer reads showed less to no signs of underestimation (Fig. S3B).

Both the dose escalation as well as progressive time course data sets were generated using the TruSeq protocol and sequenced with 2x 76bp paired end reads. Indeed, in accordance with the simulated data, the estimated 4sU incorporation frequency did not change significantly for these data sets (Fig. S4, S5), and the effect on short-lived RNA was still clearly visible after rescue (Fig. S6, S7).

We concluded that even though improved read mapping can fully mitigate the effect of reduced mappability of T-to-C conversion reads, short-lived RNA still appears to be downregulated with high 4sU concentrations at long labelling times.

### Labelled RNA is underrepresented by a constant factor

We hypothesized that a global and unspecific underrepresentation of labelled RNA in the sequencing libraries is responsible for the observed downregulation of short-lived RNAs and that gene-specific differences in RNA half-lives can explain gene-specific differences in downregulation. In this case, in each sample the same fraction of labelled RNA is missing for each gene. To test this hypothesis, we devised an algorithm to estimate this percentage of 4sU dropout and used this parameter to scale up the estimated newly synthesized RNA per gene (Fig. 6A).

**Figure 6.**
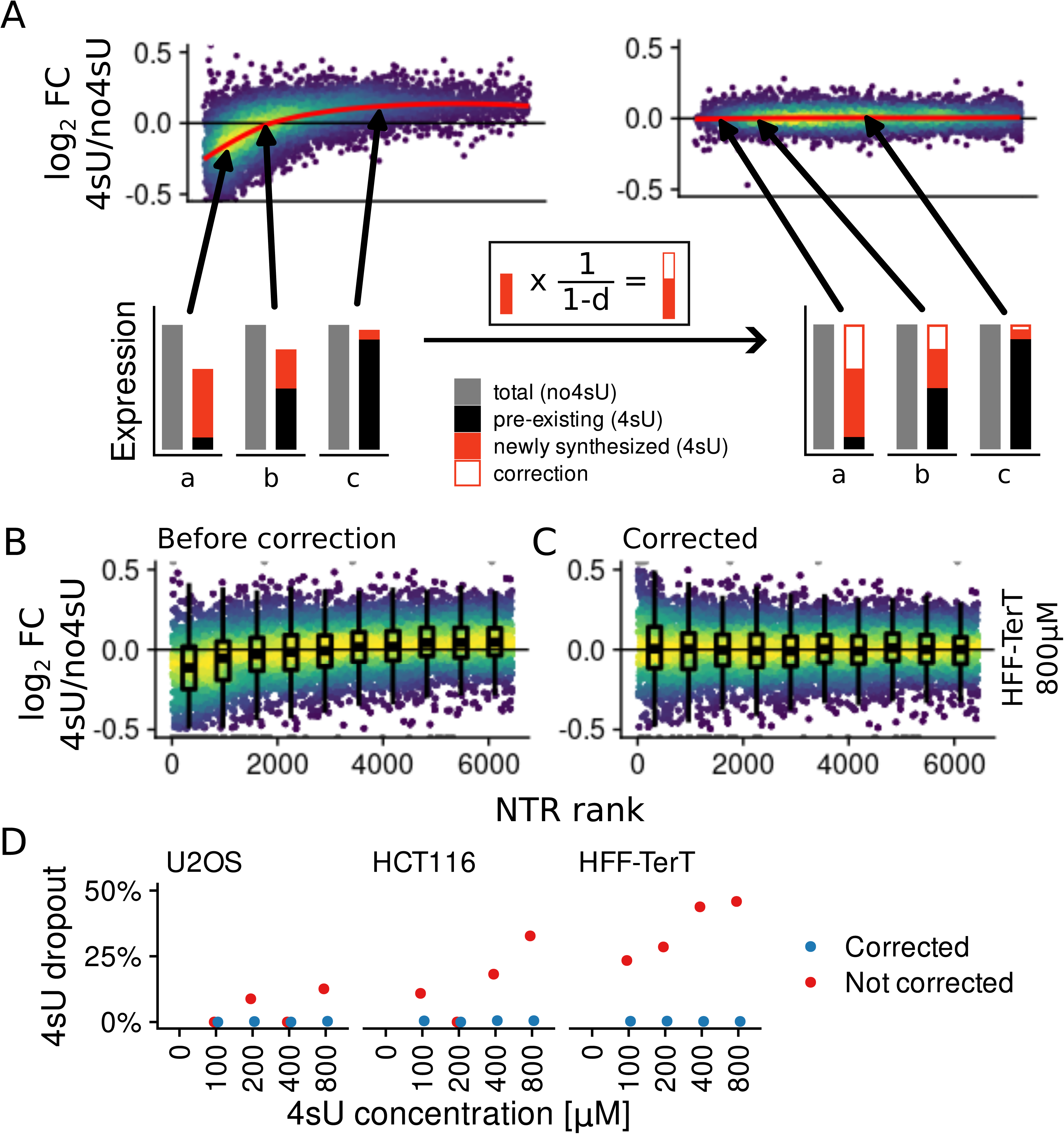
Correction of 4sU dropout by scaling labelled RNA. (A) Correction of three example genes affected by 4sU dropout with short (a), medium (b) and long RNA half-life (c). A dropout factor ‘d’ is calculated and subsequently expression of labelled RNA is multiplied by 1/1-d. (B,C) 4sU dropout rank plots of the 800 µM HFF-TerT sample before and after correction. The x axis here is the rank of the new-to-total RNA ratio. Boxplots showing the log2 fold changes of the 4sU labelled sample vs unlabelled control are overlayed for 10 equisized bins along the x axis.(D) Comparison of 4sU dropout percentage before and after correction in all three cell lines and for all 4sU concentrations.

We estimated 4sU dropout to minimize the absolute correlation of the log_2_ fold change of the 4sU treated sample vs the corresponding 4sU naïve sample (4sU vs no4sU) against the new-to-total RNA ratio per gene. Before this correction, this correlation was strong and highly significant for the 800 μM HFF-TerT sample (Fig. 6B, Spearman’s ρ=0.29, p<2.2x10, asymptotic t test). After correction, the correlation vanished (Fig. 6C, Spearman’s ρ=0, p=0.91, asymptotic t test). Importantly, we did not observe any signs of a non-monotonic correlations after correction: The distributions of the 4sU vs no4sU log_2_ fold change for 10 equisized bins along the NTRs were indistinguishable (p=0.11, Kruskall- Wallis-test). This result suggests that scaling by the percentage of transcriptional loss completely removed the observed effect of preferential downregulation of short-lived RNAs.

The 4sU dropout percentage cannot only be used to correct for this effect, but also is a convenient way to quantify the extent of this effect per sample as an alternative to visually inspecting the corresponding 4sU dropout plots. Indeed, the dropout values for the 4sU dose escalation experiments mirrored our visual impression (Fig. 6D): For HFF-TerTs, the 4sU dropout rose to >40% at 800 μM and was lower for all other samples. For the two cancer cell lines, only the 800 μM sample of HCT116 was above 30%. In summary, the 4sU dropout percentage can be used as a statistic to quantify preferential downregulation of short-lived RNA and to correct for it.

### 4sU dropout scaling mitigates biased expression estimates

To further investigate whether scaling using the 4sU dropout percentage can mitigate the effect of downregulation of short-lived RNAs, we analysed our progressive labelling time course data set. The transcriptional loss was remarkably consistent among replicates and increased steadily and almost linearly with longer labelling time up to a value of 31.6% and 33.5% for the two replicates with 2h labelling (Fig. 7A). Again, scaling labelled RNA based on the 4sU dropout percentage corrected for the downregulation effect without clear signs of non-monotonic correlations (Fig. 7B-C).

**Figure 7.**
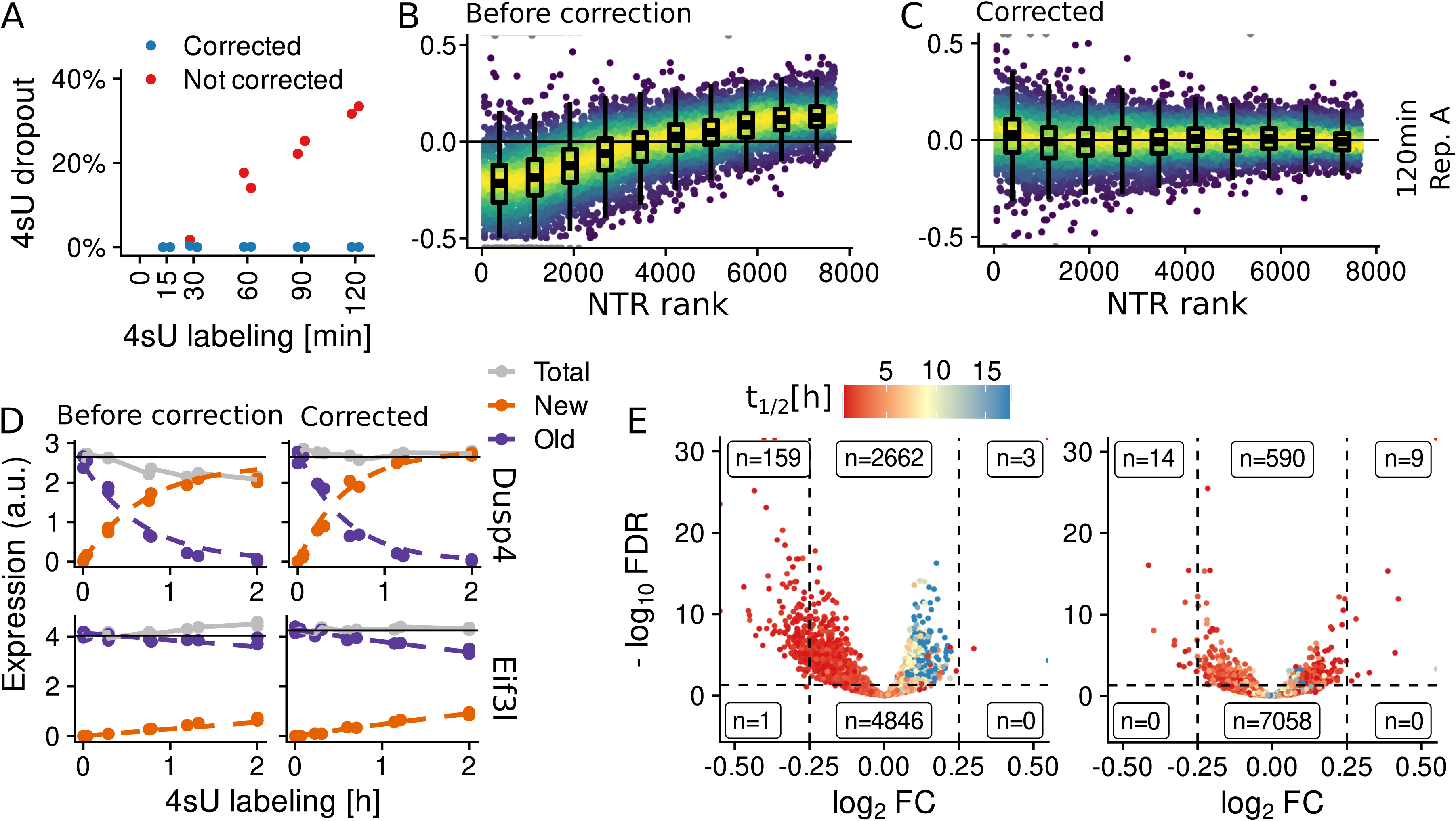
Evaluation of 4sU dropout correction in progressive labelling data. (A) Comparison of the 4sU dropout percentage before and after correction in progressive labelling data over all time points in replicate A. (B,C) 4sU dropout rank plots of the 120 min sample in replicate A before and after correction with overlayed boxplots as in Figure 6B-C(D) Old, new and total gene expression of Dusp4 and Eif3l over 2 hours of labelling before (left) and after (right) correction. The kinetic model fits are indicated as dashed lines. (E) Vulcano plot of genes showing an upwards or downwards trend on total RNA level before (left) and after correction (right). The y axis shows -log_10_ of the DESeq2 P value (likelihood ratio test comparing a model with the 4sU labelling time as independent variable vs a model with intercept only) adjusted for multiple testing (Benjamini-Hochberg; FDR, false discovery rate). The numbers of genes above and below 5% FDR and log2 fold changes of > 0.25 are indicated. Gene half-lives are represented by color.

4sU dropout does not only bias estimates of the NTRs, but also has profound effects for normalization across samples with distinct labelling times, e.g. for progressive labelling time courses. This is because under 4sU dropout, the fundamental assumption of no global changes in gene expression that most normalization methods make is violated. Indeed, normalization resulted in downward trends along the labelling time for short-lived RNAs, and in upwards trends for long-lived RNAs (Fig. 7D-E). For example, the levels of the short-lived RNA of Dusp4 declined at a rate of 12% per hour after size factor normalization (27) whereas the levels of the long-lived RNA of Eif3l increased at a rate of 5% per hour (Fig. 7D). Both trends were fully corrected by 4sU dropout scaling (Fig. 7D). Of note, these downwards and upwards trends due to normalization also bias half-life estimates: Upon correction, the estimated half-lives changed from 28.7 min (0.95% CI: 25.5 - 32.8 min) to 23.0 min (0.95% CI: 20.6 – 26.0 min) for Dusp4 and from 9:39h (0.95% CI: 7:56 – 12:19h) to 5:51h (0.95% CI: 5:22 - 6:25h) for Eif3l. Globally, n=162 genes show a strong and significant upwards or downwards trend without correction (absolute log2 fold change per hour > 0.25, P value < 5%, likelihood ratio test, Benjamini-Hochberg adjusted for multiple testing, Fig. 7E), and only n=23 remain after transcriptional loss factor scaling (Fig. 7E). In summary, scaling by the 4sU dropout percentage removed global 4sU induced effects on expression estimates that occur with high 4sU concentrations and long periods of labelling.

### Impaired reverse transcription results in 4sU dropout

In principle, extensive 4sU dropout observed in our progressive labelling data set could be due to a direct or indirect effect of 4sU on RNA metabolism in the living cells, or because labelled RNA is underrepresented in the sequencing library due to diminished reverse transcription efficiency of 4sU containing RNA. To test this hypothesis, we determined the RNA fragments that were reverse transcribed from the paired-end sequencing data for all samples for n=105 genes that had an RNA half-life of <30 min, were strongly expressed (>10 TPM) and had an estimated major isoform percentage of >90%. Interestingly, RNA fragments across all 105 genes that were sequenced from cells that were treated with 4sU for 2h had significantly lower U or 4sU content than RNA fragments sequenced from 4sU naïve cells (p<2.2x10, Wilcoxon test, Fig. 8A-B). Lower U or 4sU content was consistent for both replicates and gradually decreased with the labelling time (Fig. 8C). This was not due to issues with read mappability since reads were mapped using grand-Correct, and we also observed the same differences in nucleotide content for the not sequenced parts of the RNA fragments in between the read pair (Fig. S8). Notably, by counting di- and trinucleotides, we found that underrepresentation of U or 4sU in the RNA fragments in 4sU labelled samples depended on the sequence context with neighbouring U (or 4sU), A or G nucleotides resulting in stronger underrepresentation (Fig. 8D and S8). Taken together, this suggests that reverse transcription efficiency of iodoacetaminde-converted 4sU nucleotides is impaired.

**Figure 8.**
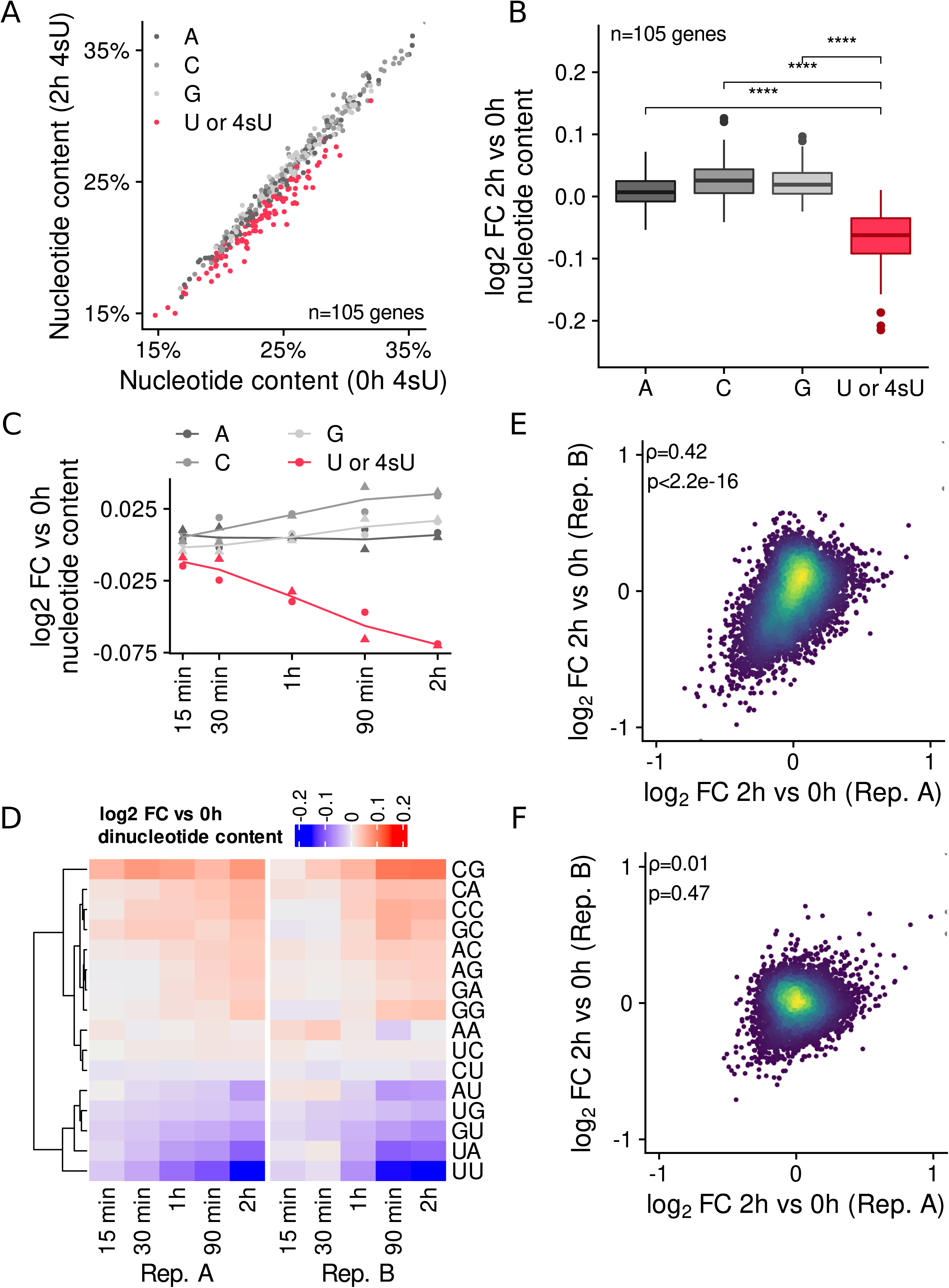
4sU incorporation reduces reverse transcription efficiency. (A) Scatter plot of nucleotide content in sequenced RNA fragments in 4sU naïve sample vs. 2 hours 4sU labelling. n=105 genes with an RNA half-life <30 min, >10TPM and an estimated major isoform percentage of >90% are shown. (B) Boxplots showing the log2 fold changes for all n=105 genes of nucleotide content in 2 hours 4sU labelling vs. 4sU naïve sample for all nucleotides. P values (<2.2x10^-16^, Wilcoxon test) are indicated. (C) Line plots showing the average log2 fold changes across the n=105 genes of nucleotide content over all labelling times vs. 4sU naïve sample and both replicates. (D) Heatmaps showing the average log2 fold changes across all n=105 genes for all dinucleotides over all labelling times vs. 4sU naïve sample in both replicates. (E,F) Scatter plot of log2 fold changes for 2 hours labelling vs. 4sU naïve sample in replicate A against log2 fold changes in replicate B before (E) and after (F) correction. The Spearman correlation coefficient and associated P values (asymptotic t test) are indicated.

These findings indicate that inefficient reverse transcription of 4sU is responsible for the bias in the expression estimates. In this case, 4sU dropout is a global and random effect that affects all fragments from labelled RNAs to roughly the same extent. Any other covariate that correlates with the 4sU vs no4sU log_2_ fold change would also result in a correlation of the 4sU vs no4sU log_2_ fold change among replicates. Indeed, these fold changes of the uncorrected 2h replicates were strongly correlated (Spearman’s ρ=0.42, p<2.2x10^-16^, asymptotic t test, Fig 8E). However, after scaling this correlation disappeared completely (Spearman’s ρ=0.01, p=0.47, asymptotic t test, Fig. 8F). Thus, any other additional factor resulting in differences between 4sU treated and 4sU naïve samples was minor in comparison to biological variability among samples. In summary, these findings indicate that scaling by the 4sU dropout percentage can fully correct for bias in expression estimates due to excessive 4sU labelling.

## DISCUSSION

Nucleotide conversion RNA-seq requires high 4sU concentrations: The 4sU concentration correlates with the 4sU incorporation frequency in labelled RNA, which in turn determines how many reads originating from labelled RNA carry a T-to-C mismatch. For the data from Ref. (1), we estimated an incorporation frequency in labelled RNA of 2% (11). Based on binomial statistics, with the 50bp single end reads that were used in this study, 75% of all reads originating from labelled RNA are expected to carry no T-to-C mismatch (10). With higher concentrations resulting in an incorporation frequency of 10%, about 80% would carry at least one T-to-C mismatch. Statistical approaches such as GRAND-SLAM can deal with such missing observations, but also benefit substantially from higher incorporation frequencies (11). In addition, the accuracy of half-life estimates drop severely when the labelling time is much shorter than the RNA half-life (12). Thus, in addition to high concentrations, long periods of labelling are required for accurately estimating the whole spectrum of RNA half-lives for mammalian genes. However, excessive labelling with 4sU reduces cell viability (1) and has been shown to affect rRNA processing (13). In addition to these biological effects, we show here that excessive labelling also affects sequencing data due to reduced reverse transcription efficiency and mappability of reads with many mismatches.

String matching allowing for mismatches is computationally a much harder problem than exact string matching (28). Therefore, all available read mapping tools use a two-step approach to quickly map reads: First, using a data structure for exact string matching and some heuristics to allow for mismatches, candidate mapping positions are identified. Second, the candidate positions are then filtered according to user-defined criteria such as the number of maximal mismatches. To improve read mapping for T-to-C mismatches, two different strategies have been proposed: HISAT-3N (26) operates on a genome with reduced, three-letter alphabet, and SLAM-dunk (25) uses adapted criteria that do not penalize T-to-C mismatches in the filtering step. Thus, HISAT-3N is aware of 4sU induced conversions for both candidate generation and filtering, while SLAM-dunk considers conversions only for filtering. Both strategies have disadvantages that could be observed for our simulated reads: Mapping with reduced alphabets generates more multi-mappers, whereas conversion aware filtering misses true mapping locations with increasing numbers of mismatches. Our grand-Rescue approach is also based on a reduced alphabet, but we mitigate the effect of multi- mappers by only mapping previously unmappable reads against a three-letter pseudo-transcriptome. More importantly, grand-Rescue is in principle agnostic of the underlying read mapping tool that also has major impact on the overall performance.

Reduced reverse transcription (RT) efficiency of 4sU containing RNA, as described recently (14), can also result in 4sU dropout in nucleotide conversion RNA-seq. With reduced RT efficiency, the same percentage of labelled RNA is missing for all genes in the sequencing library. Here, we showed that this percentage can be estimated from data and can be used to computationally remove 4sU dropout. The effect of RT efficiency depends on the library preparation protocol. All data here were generated using the Illumina TruSeq stranded mRNA kit. In this protocol, purified polyadenylated mRNA is randomly fragmented, and cDNA from these fragments is made from random hexamer primers. If each 4sU nucleotide in the RNA fragment reduces the processivity of the reverse transcriptase, 4sU dropout correlates with the number of 4sU nucleotides in the RNA fragment. In principle, the uridine content of mRNAs could be used as a covariate when correcting 4sU dropout. For several reasons, we here resided with a more parsimonious model: First, it is impossible to evaluate, whether including uridine content would provide improved quantifications. Second, we expect the influence of uridine content to even out when RNA fragments over full length mRNAs are sequenced. Third, we also observed dependence on surrounding nucleotides, which would suggest even more complex models. Finally, our data indicate that also with a simpler model, the influence of RT efficiency could be removed from the data.

Implementations of the two methods introduced here are available as part of our GRAND-SLAM/grandR pipeline. grand-Rescue is a stand-alone program that is integrated as an additional step into the pipeline. Its only input is a single bam file containing also unmapped reads, and it generates a new bam file containing the rescued read mappings in addition to previously mapped reads. Thus, it can be integrated into any existing pipeline as an additional step. Computation of the 4sU dropout percentage and the scaling approach to correct for dropout are implemented as functions in our grandR package (12), and we provide a vignette to showcase the usage of these functions. In principle, computing the 4sU dropout percentage of a sample that has been labelled with 4sU requires an otherwise biologically equivalent control sample without 4sU labelling as reference, or a reference sample without a global change in RNA synthesis or stability.

Here, we report that high concentrations of 4sU or prolonged labelling resulted in an apparent downregulation of short-lived RNAs, which can have profound impact on results when staying unnoticed. We therefore advocate that checking for this effect is a mandatory part of quality control for nucleotide conversion RNA-seq. If such quantification bias is observed, it is important to investigate its causes. Technical issues such as reduced read mappability for labelled RNA or inefficient reverse transcription of labelled RNA can result in 4sU dropout und therefore in apparent downregulation of short-lived RNA. Indeed, for the strong effect observed after 2h labelling in NIH- 3T3 cells with 800μM 4sU, we provide evidence that reverse transcription efficiency played a major role. It is not unlikely, that this technical issue might be the only cause for 4sU dropout in this experiment, since after correction by our scaling approach, no quantification bias was observed anymore, and the correlation of log2 fold changes between replicates disappeared completely. However, downregulation of short-lived RNA can also be a sign of 4sU affecting the living cells biologically. If such an effect of 4sU on RNA metabolism cannot be excluded, all obtained results can be misleading and must be interpreted with care.

## DATA AVAILABILITY

Raw data generated here have been deposited at GEO under accession numbers GSE229504 (Increasing concentrations), and GSE229506 (Progressive labelling).

All processed data (GRAND-SLAM outputs) are available on zenodo:

- https://doi.org/10.5281/zenodo.7805929 (progressive labelling, Ref. 4)
- https://doi.org/10.5281/zenodo.7753460 (Increasing concentrations)
- https://doi.org/10.5281/zenodo.7760483 (QuantSeq (17)),
- https://doi.org/10.5281/zenodo.7760437 (Illumina TruSeq (16))

R notebooks and data files for generating all figures are available on zenodo (https://doi.org/10.5281/zenodo.7842413).

## AUTHOR CONTRIBUTIONS

Kevin Berg: Formal Analysis, Methodology, Software, Visualization, Writing-original draft, Writing-review & editing. Manivel Lodha: Investigation. Yiliam Cruz Garcia: Investigation. Thomas Hennig: Supervision. Elmar Wolf: Supervision. Bhupesh K Prusty: Investigation, Supervision. Florian Erhard: Conceptualization, Formal Analysis, Funding acquisition, Methodology, Software, Supervision, Visualization, Writing-original draft, Writing-review & editing

---

**Figure S1.**
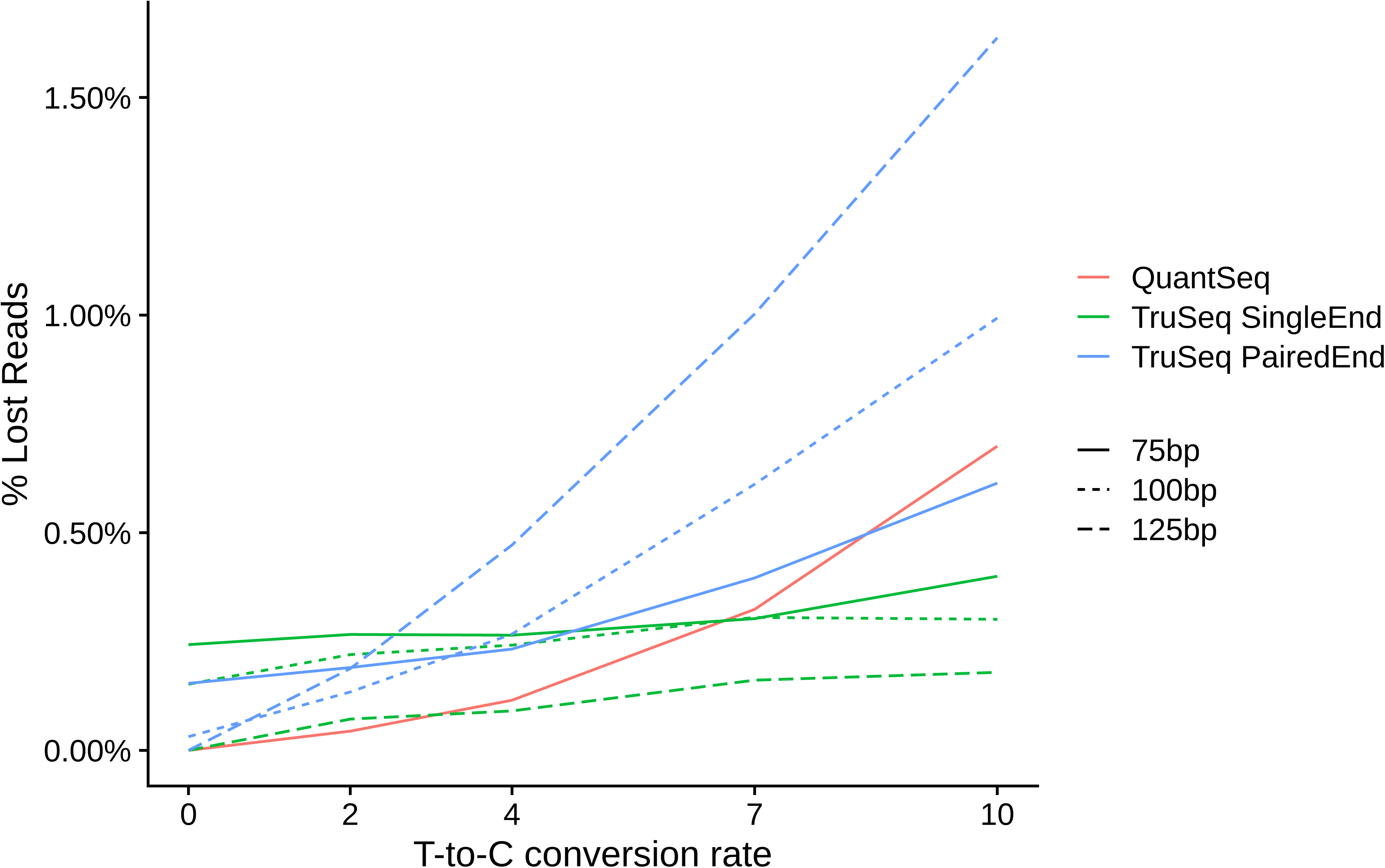
Influence of library preparation technique and read lengths on read mappability. Line plots showing the percentage of lost reads over increasing mismatch rates in QuantSeq, single and paired end TruSeq and read lengths of 75 bp, 100 bp and 125 bp.

**Figure S2.**
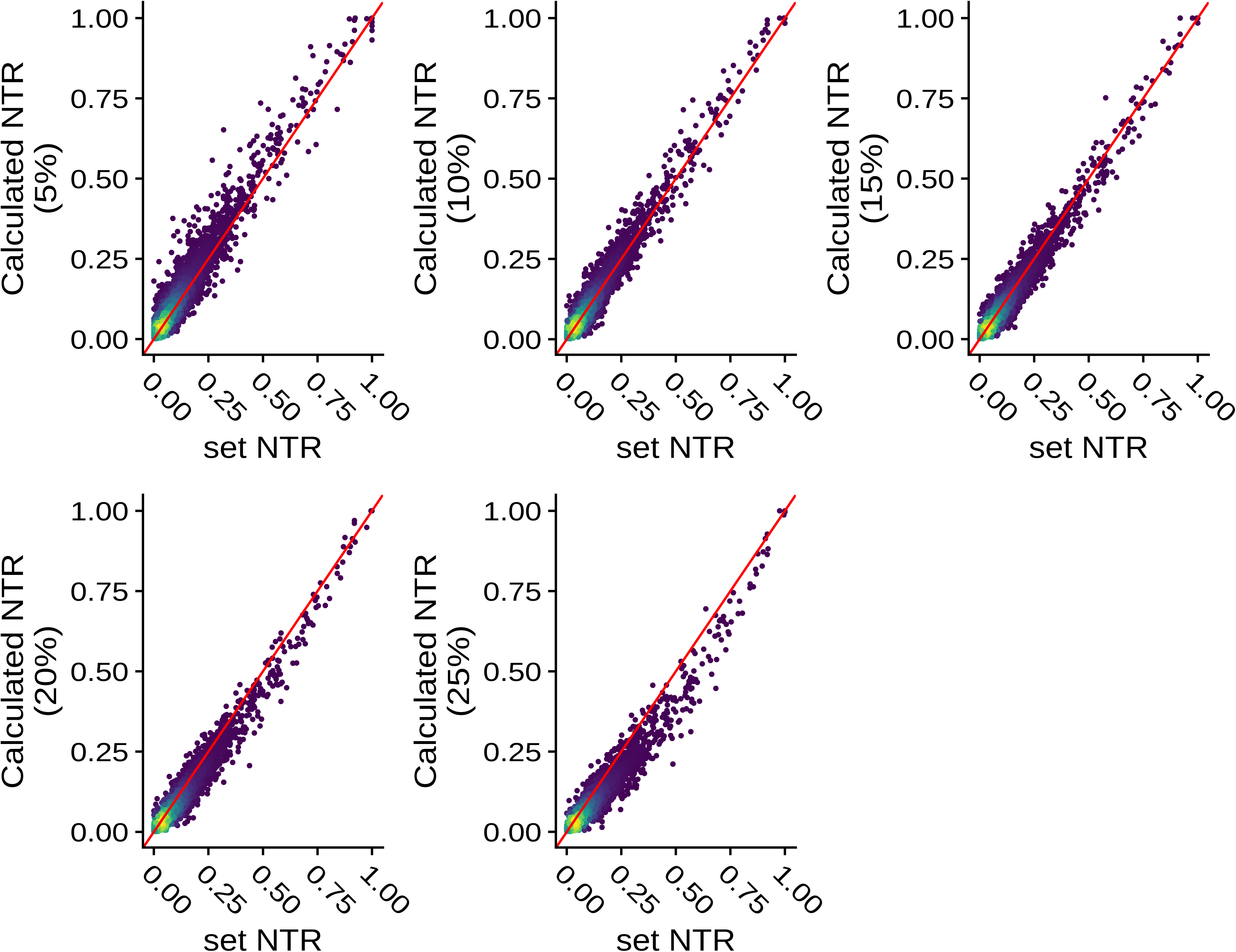
Estimation of gene-wise NTRs over increasing conversion rates. The NTRs defined by the simulation, are scattered against the estimated NTRs after simulation with conversion rates from 5% to 25%.

**Figure S3.**
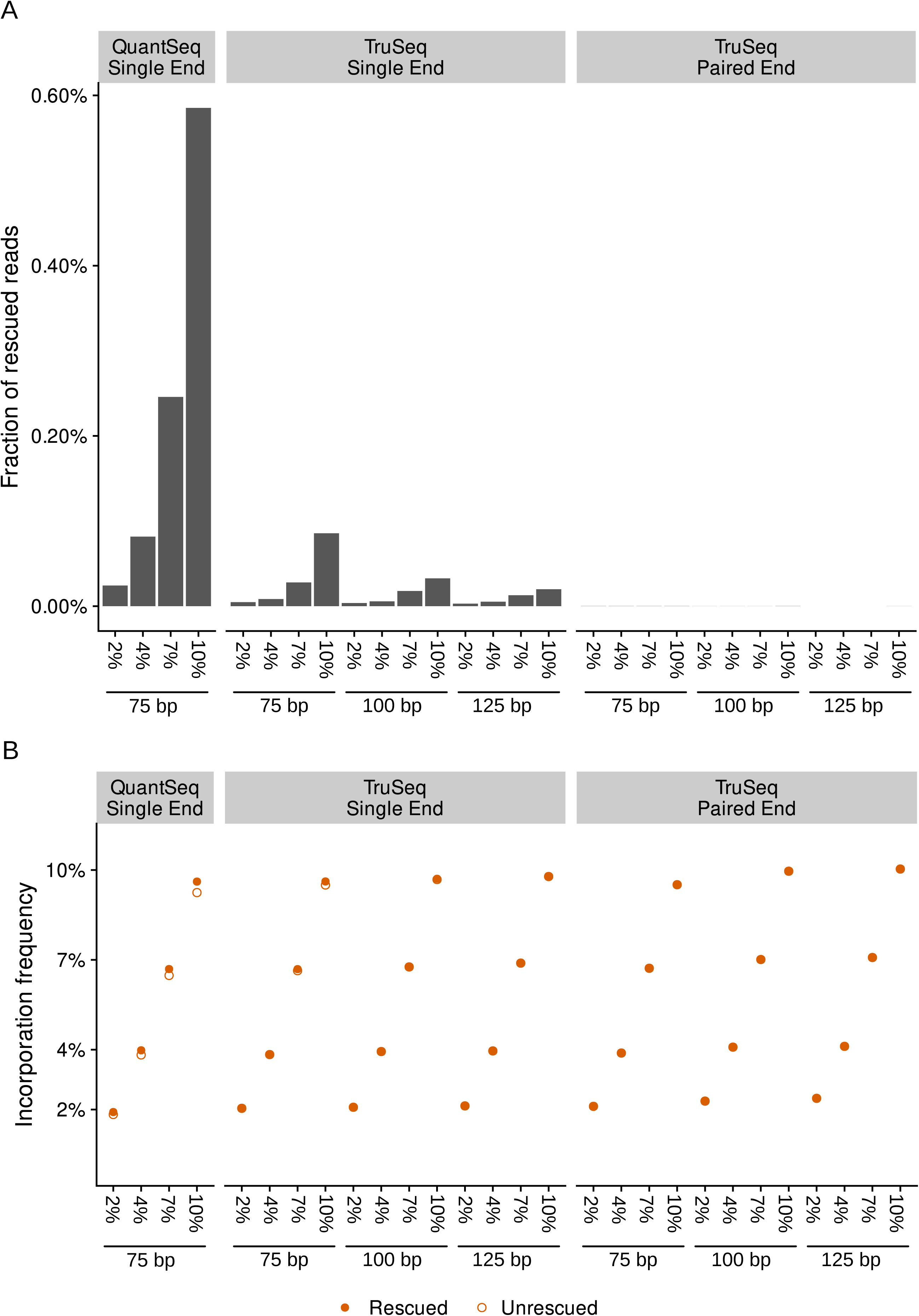
Effectiveness of grand-Rescue in simulations by library preparation method and read lengths. (A) Fraction of rescued reads by grandRescue relative to all reads per sample for QuantSeq, TruSeq (single and paired end), read lengths of 75 bp, 100 bp, 125bp and for conversion rates from 2% to 10%. (B) GRAND-SLAM estimates of 4sU incorporation rates before (empty dots) and after (solid dots) rescue for QuantSeq, TruSeq (single and paired end), read lengths of 75 bp, 100 bp, 125bp and for conversion rates from 2% to 10%.

**Figure S4.**
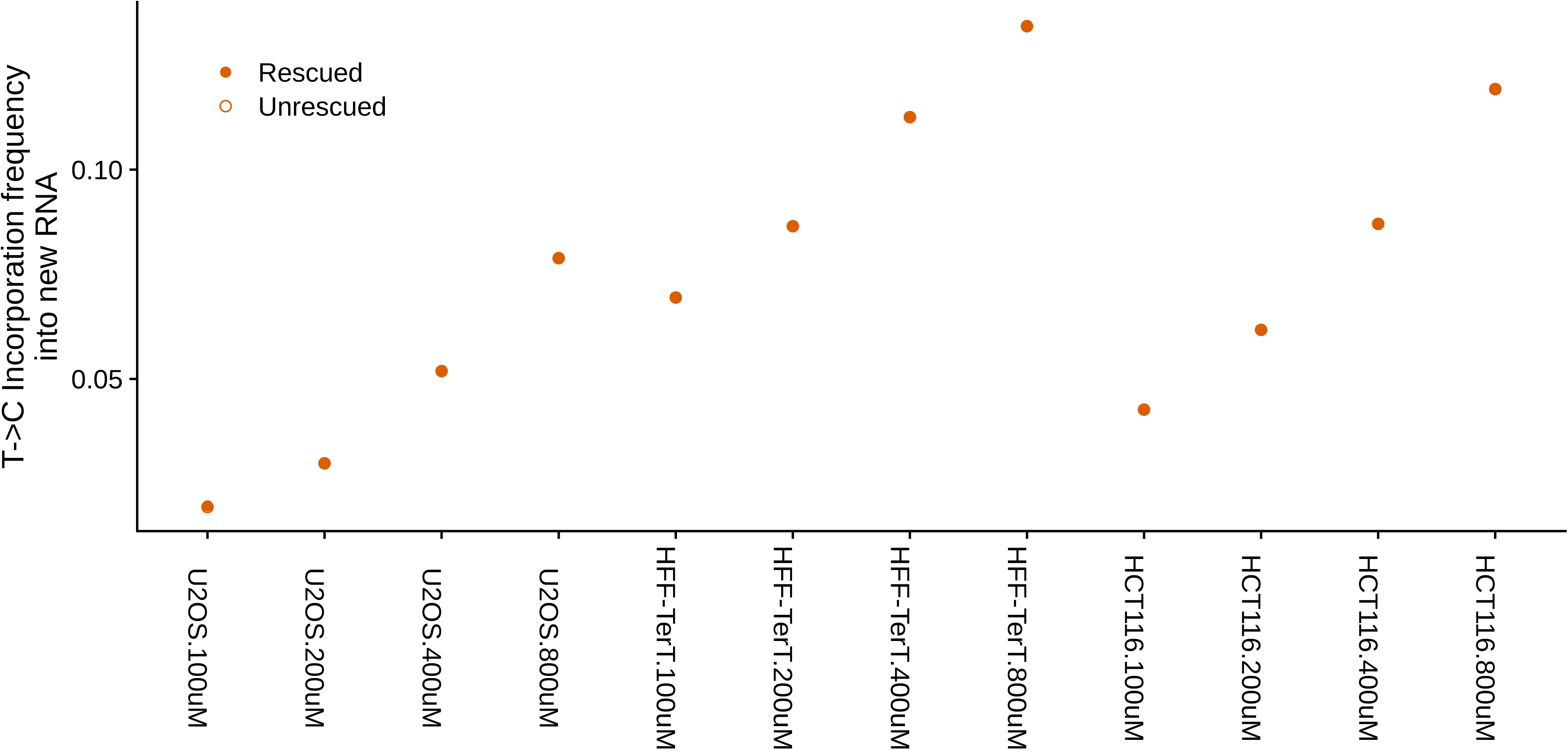
Estimated incorporation frequencies in increasing concentration samples. GRAND-SLAM estimates of incorporation frequencies before and after rescue for all the cell lines and labelling concentrations of 100 µM, 200 µM, 400 µM and 800 µM.

**Figure S5.**
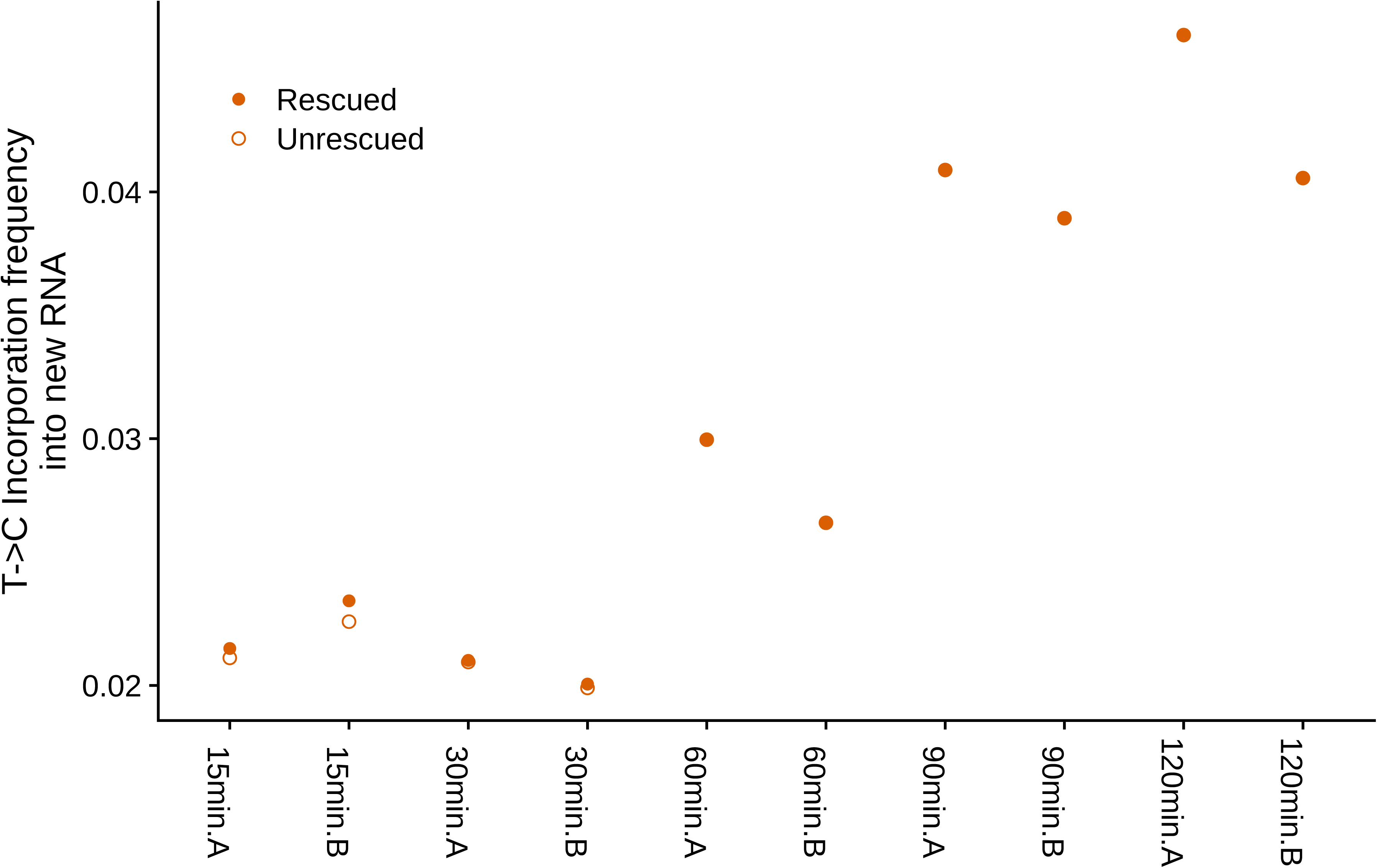
Estimated incorporation frequencies in progressive labelling samples. GRAND-SLAM estimates of incorporation frequencies before and after rescue in both replicates for 15 min, 30 min, 60 min, 90 min and 120 min of labelling.

**Figure S6.**
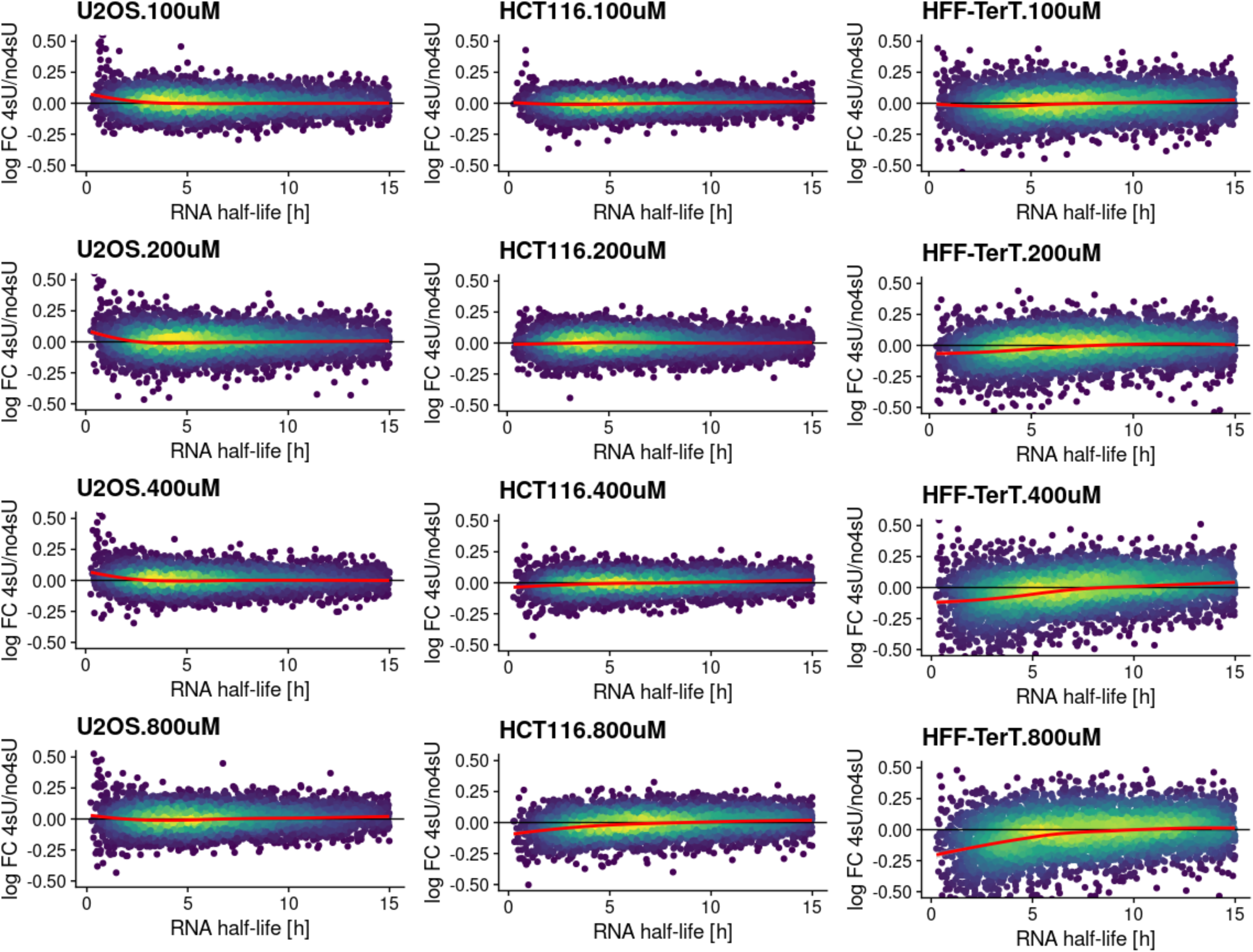
4sU dropout plots of n=6,454 genes for all 4sU samples vs. corresponding 4sU naïve samples for all three cell lines and all 4sU concentrations. The x axis shows the RNA half-life, the y axis the median centered log2 fold change of total RNA expression for the 4sU labelled sample vs. the corresponding 4sU naïve control sample. A local polynomial regression (loess) fit is indicated in red.

**Figure S7.**
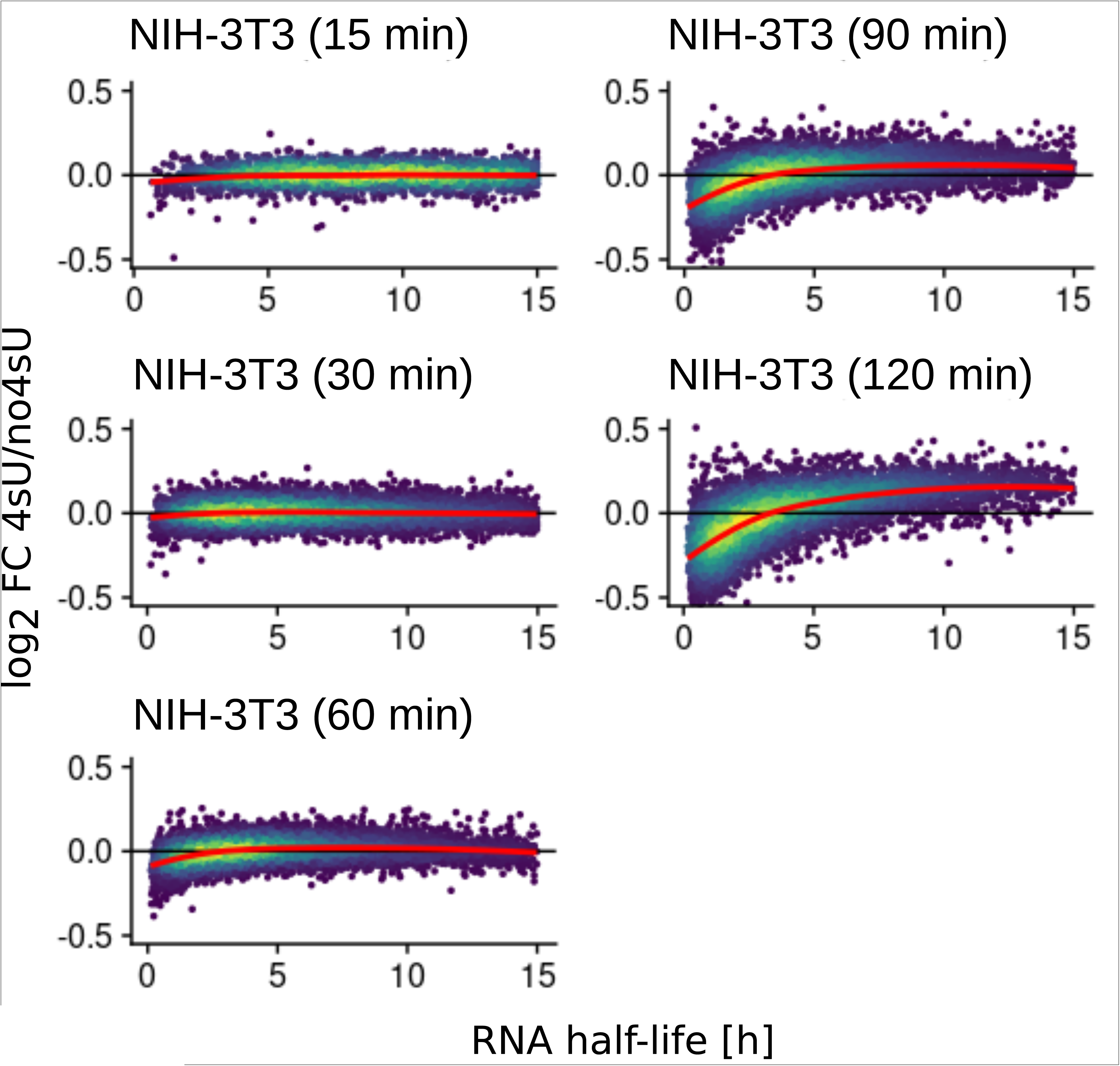
4sU dropout plots of n=9,072 genes for all time points of replicate B. The x axis shows the RNA half-life, the y axis the median centered log2 fold change of total RNA expression for the 4sU labelled sample vs. the corresponding 4sU naïve control sample. A local polynomial regression (loess) fit is indicated in red.

**Figure S8.**
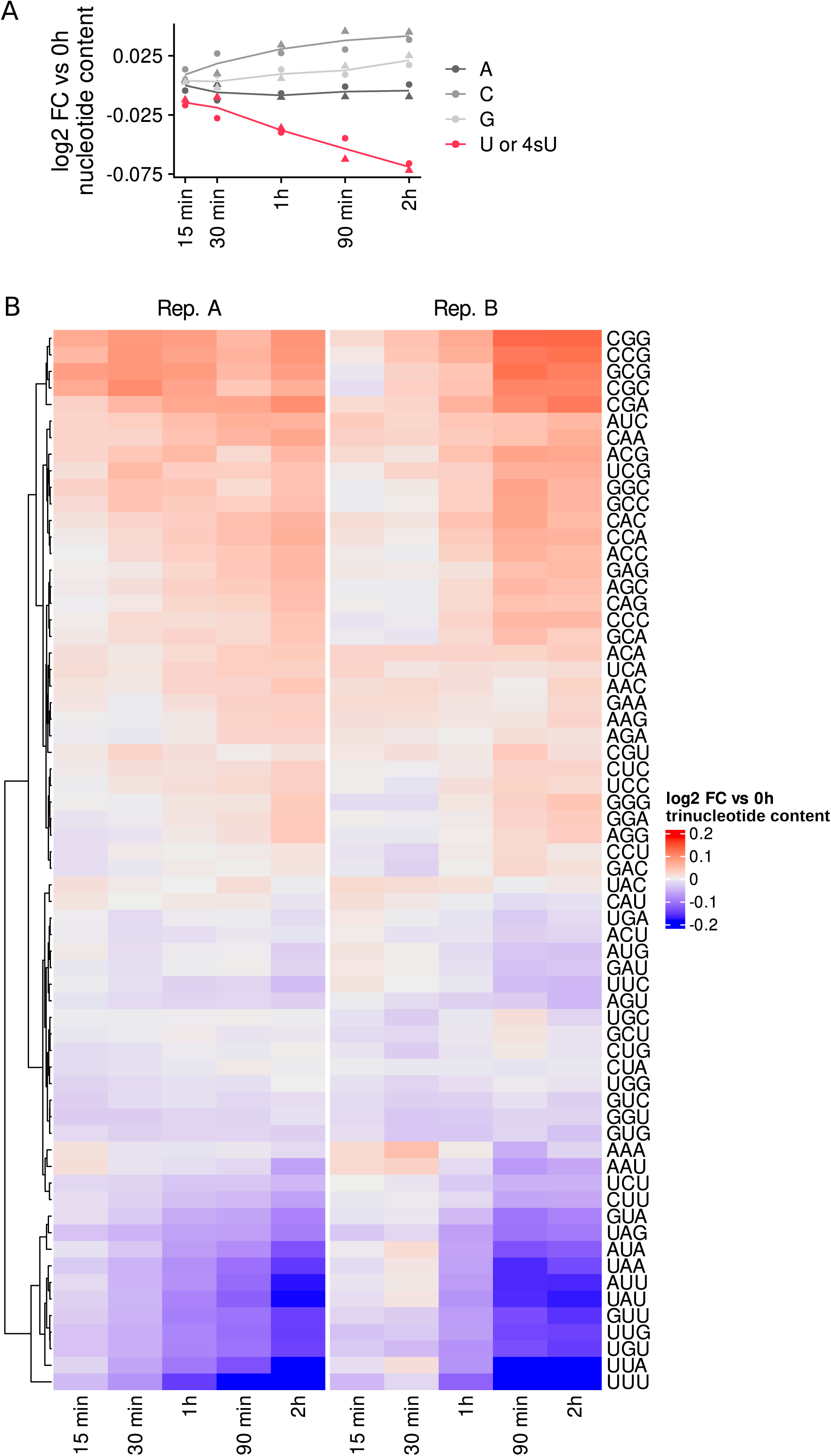
Nucleotide content in unsequenced parts and trinucleotide representation in progressive labelling data. (A) Line plots showing the log2 fold change of nucleotide content in unsequenced parts of RNA fragments in between read pairs over all labelling times vs. 4sU naïve sample in both replicates (shapes). (B) Heatmaps showing log2 fold change for all trinucleotides over all labelling times vs. 4sU naïve sample in both replicates.

## REFERENCES

1. Herzog, V.A., Reichholf, B., Neumann, T., Rescheneder, P., Bhat, P., Burkard, T.R., Wlotzka, W., Haeseler, A. von, Zuber, J. and Ameres, S.L. (2017) Thiol-linked alkylation of RNA to assess expression dynamics. Nature Methods, 14, 1198.

2. Schofield, J.A., Duffy, E.E., Kiefer, L., Sullivan, M.C. and Simon, M.D. (2018) TimeLapse-seq: adding a temporal dimension to RNA sequencing through nucleoside recoding. Nature Methods, 15, 221–225.

3. Riml, C., Amort, T., Rieder, D., Gasser, C., Lusser, A. and Micura, R. (2017) Osmium-Mediated Transformation of 4-Thiouridine to Cytidine as Key To Study RNA Dynamics by Sequencing. Angew. Chem. Int. Ed. Engl., 56, 13479–13483.

4. Narain, A., Bhandare, P., Adhikari, B., Backes, S., Eilers, M., Dölken, L., Schlosser, A., Erhard, F., Baluapuri, A. and Wolf, E. (2021) Targeted protein degradation reveals a direct role of SPT6 in RNAPII elongation and termination. Mol Cell, 10.1016/j.molcel.2021.06.016.

5. Finkel, Y., Gluck, A., Nachshon, A., Winkler, R., Fisher, T., Rozman, B., Mizrahi, O., Lubelsky, Y., Zuckerman, B., Slobodin, B., et al. (2021) SARS-CoV-2 uses a multipronged strategy to impede host protein synthesis. Nature, 594, 240– 245.

6. Erhard, F., Baptista, M.A.P., Krammer, T., Hennig, T., Lange, M., Arampatzi, P., Jürges, C.S., Theis, F.J., Saliba, A.-E. and Dölken, L. (2019) scSLAM-seq reveals core features of transcription dynamics in single cells. Nature, 571, 419–423.

7. Cao, J., Zhou, W., Steemers, F., Trapnell, C. and Shendure, J. (2020) Sci-fate characterizes the dynamics of gene expression in single cells. Nat Biotechnol, 38, 980–988.

8. Schott, J., Reitter, S., Lindner, D., Grosser, J., Bruer, M., Shenoy, A., Geiger, T., Mathes, A., Dobreva, G. and Stoecklin, G. (2021) Nascent Ribo-Seq measures ribosomal loading time and reveals kinetic impact on ribosome density. Nat Methods, 18, 1068–1074.

9. Jürges, C.S., Lodha, M., Le-Trilling, V.T.K., Bhandare, P., Wolf, E., Zimmermann, A., Trilling, M., Prusty, B., Dölken, L. and Erhard, F. (2022) Multi-omics reveals principles of gene regulation and pervasive non-productive transcription in the human cytomegalovirus genome. 10.1101/2022.01.07.472583.

10. Erhard, F., Saliba, A.-E., Lusser, A., Toussaint, C., Hennig, T., Prusty, B.K., Kirschenbaum, D., Abadie, K., Miska, E.A., Friedel, C.C., et al. (2022) Time-resolved single-cell RNA-seq using metabolic RNA labelling. Nat Rev Methods Primers, 2, 1–18.

11. Jürges, C., Dölken, L. and Erhard, F. (2018) Dissecting newly transcribed and old RNA using GRAND-SLAM. Bioinformatics, 34, i218–i226.

12. Rummel, T., Sakellaridi, L. and Erhard, F. (2022) grandR: a comprehensive package for nucleotide conversion sequencing data analysis. 10.1101/2022.09.12.507665.

13. Burger, K., Mühl, B., Kellner, M., Rohrmoser, M., Gruber-Eber, A., Windhager, L., Friedel, C.C., Dölken, L. and Eick, D. (2013) 4-thiouridine inhibits rRNA synthesis and causes a nucleolar stress response. RNA Biol, 10, 1623–1630.

14. Watson, M.J., Park, Y. and Thoreen, C.C. (2021) Roadblock-qPCR: a simple and inexpensive strategy for targeted measurements of mRNA stability. RNA, 27, 335–342.

15. Whisnant, A.W., Jürges, C.S., Hennig, T., Wyler, E., Prusty, B., Rutkowski, A.J., L’hernault, A., Djakovic, L., Göbel, M., Döring, K., et al. (2020) Integrative functional genomics decodes herpes simplex virus 1. Nat Commun, 11, 1–14.

16. Sarantopoulou, D., Tang, S.Y., Ricciotti, E., Lahens, N.F., Lekkas, D., Schug, J., Guo, X.S., Paschos, G.K., FitzGerald, G.A., Pack, A.I., et al. (2019) Comparative evaluation of RNA-Seq library preparation methods for strand-specificity and low input. Sci Rep, 9, 13477.

17. Lee, J.W., Stone, M.L., Porrett, P.M., Thomas, S.K., Komar, C.A., Li, J.H., Delman, D., Graham, K., Gladney, W.L., Hua, X., et al. (2019) Hepatocytes direct the formation of a pro-metastatic niche in the liver. Nature, 567, 249–252.

18. Erhard, F. (2018) Estimating pseudocounts and fold changes for digital expression measurements. Bioinformatics, 34, 4054–4063.

19. Dölken, L., Ruzsics, Z., Rädle, B., Friedel, C.C., Zimmer, R., Mages, J., Hoffmann, R., Dickinson, P., Forster, T., Ghazal, P., et al. (2008) High-resolution gene expression profiling for simultaneous kinetic parameter analysis of RNA synthesis and decay. RNA, 14, 1959–1972.

20. Smalec, B.M., Ietswaart, R., Choquet, K., McShane, E., West, E.R. and Churchman, L.S. (2022) Genome-wide quantification of RNA flow across subcellular compartments reveals determinants of the mammalian transcript life cycle. 10.1101/2022.08.21.504696.

21. Dobin, A., Davis, C.A., Schlesinger, F., Drenkow, J., Zaleski, C., Jha, S., Batut, P., Chaisson, M. and Gingeras, T.R. (2013) STAR: ultrafast universal RNA-seq aligner. Bioinformatics, 29, 15–21.

22. Krueger, F. and Andrews, S.R. (2011) Bismark: a flexible aligner and methylation caller for Bisulfite-Seq applications. Bioinformatics, 27, 1571–1572.

23. Baruzzo, G., Hayer, K.E., Kim, E.J., Di Camillo, B., FitzGerald, G.A. and Grant, G.R. (2017) Simulation-based comprehensive benchmarking of RNA-seq aligners. Nat Methods, 14, 135–139.

24. Kim, D., Paggi, J.M., Park, C., Bennett, C. and Salzberg, S.L. (2019) Graph-based genome alignment and genotyping with HISAT2 and HISAT-genotype. Nat Biotechnol, 37, 907–915.

25. Neumann, T., Herzog, V.A., Muhar, M., von Haeseler, A., Zuber, J., Ameres, S.L. and Rescheneder, P. (2019) Quantification of experimentally induced nucleotide conversions in high-throughput sequencing datasets. BMC Bioinformatics, 20, 258.

26. Zhang, Y., Park, C., Bennett, C., Thornton, M. and Kim, D. (2021) Rapid and accurate alignment of nucleotide conversion sequencing reads with HISAT-3N. Genome Res., 10.1101/gr.275193.120.

27. Love, M.I., Huber, W. and Anders, S. (2014) Moderated estimation of fold change and dispersion for RNA-seq data with DESeq2. Genome Biology, 15, 550.

28. Cohen-Addad, V., Feuilloley, L. and Starikovskaya, T. (2018) Lower bounds for text indexing with mismatches and differences. 10.48550/arXiv.1812.09120.

